# SETD6 Mediates Selective Interaction and Genomic Occupancy of BRD4 and MITF in Melanoma cells

**DOI:** 10.1101/2025.06.29.662016

**Authors:** Tzofit Elbaz Biton, Michal Feldman, Tomer Davidy, Nili Tickotsky Moskovitz, Liron Levin, Daniel Sevilla, Colin R. Goding, Emily Bernstein, Dan Levy

**Author notes:** Correspondence should be addressed to D.L.

## Abstract

Aberrant transcriptional programs mediate malignant transformation of melanoma, the most aggressive form of skin cancer. The lysine methyltransferase SETD6 has been implicated in regulating transcription, cell adhesion, migration, and other processes in various cancers, however its role in melanoma remains unexplored. We recently reported that SETD6 mono-methylates the BRD4 at K99 to selectively regulate transcription of genes involved in mRNA translation. Here, we observed that BRD4 methylation at K99 by SETD6 occurs in melanoma cells. Knockout of SETD6 or a point mutation at BRD4-K99 disrupts BRD4 genomic occupancy. In addition, we show that SETD6 interacts with MITF, a master transcription factor in melanocytes and melanoma, and influences the genomic distribution of MITF. Mechanistically, we uncover a novel chromatin-localized interaction between BRD4 and MITF in melanoma. Our data suggest that BRD4 binds MITF in melanoma cells and that this interaction is dependent on both SETD6-mediated methylation of BRD4 and MITF acetylation. This chromatin complex plays a pivotal role in selective recruitment of BRD4 and MITF to different genomic loci in melanoma cells.

## Introduction

Melanoma, a neoplasm of melanocytic origin, is the most severe human skin cancer(1). Malignant transformation in melanoma is caused by mutations in genes which are responsible for cell proliferation and apoptosis, epigenetic changes, changes of adhesion ability, or production of autocrine growth factors. These aspects disturb the signal transduction pathways in melanocytes, the skin cells producing the pigment melanin (2). Recent studies have revealed a complex involvement of epigenetic mechanisms in the regulation of transcriptional programs in melanoma, including methylation, chromatin modification and remodeling, as well as the diverse activities of non-coding RNAs and transcription factors (TFs) activity(3). The microphthalmia-associated transcription factor (MITF) is a master regulator of melanocyte development and controls many aspects of melanocyte and melanoma biology including the cell cycle, metabolism, DNA damage repair, survival, differentiation and proliferation(4). Further, MITF is amplified in a fraction of human melanomas (1) and has been termed a lineage survival oncogene(5).

The transcription regulator BRD4 is a member of the bromodomain and extra-terminal domain (BET) protein family. BRD4 contains two conserved bromodomains that specifically recognize acetylated lysine residues on histone and non-histone proteins(6,7). In recent years, many studies have demonstrated a notable therapeutic potential of BET inhibitors, which target BRD family members and displace them from chromatin(8,9). In melanoma, BRD4 is overexpressed and essential for tumor growth *in vivo*(10) and interacts with MITF to regulate the expression of genes important for melanin synthesis in melanocytes(11). Given this role of BRD4 in melanocyte biology, we hypothesized that SETD6-mediated methylation of BRD4 would modulate the transcriptional program in melanoma cells.

Post-translational modifications (PTMs) contribute to changing protein properties, transducing cellular signals, and regulating protein-protein interactions, thus serving as an important mechanism for the regulation of many signaling pathways and biological processes(12). Several studies have demonstrated the importance of lysine methylation for regulating epigenetic processes and fundamental cellular signaling pathways via methylation of non-histone proteins(13–15). As part of its structural characteristics, lysine can be mono-, di- or trimethylated, whereby S-adenosyl-L-methionine (SAM/AdoMet) serves as the methyl donor for this PTM. Lysine methylation is a dynamic modification that is written by protein lysine methyltransferases (PKMTs) and removed by lysine demethylases (KDMs)(16). There are approximately 50 members of the PKMT family, of which most contain a conserved Su(var), Enhancer of zeste, trithorax (SET) domain which is responsible for the enzymatic activity(17). SET domain-containing protein 6 (SETD6) is a mono-methyltransferase containing the catalytic SET domain and a Rubisco substrate-binding domain which mediates protein-protein interactions(17,18). The enzymatic activity of SETD6 was shown to be involved in the regulation of diverse processes including transcription, cell adhesion, and migration, among several others(17–29). However, the role of SETD6 in melanoma and the pathways through which it might act remain unclear. We recently showed that SETD6 functions as a molecular switch which methylates chromatin-bound BRD4 at lysine-99 (K99). Consequently, BRD4 methylation affects the recruitment of the E2F1 transcription factor to genes involved in mRNA translation in breast cancer cells(28).

Here we show that SETD6 binds and methylates BRD4 at K99 in melanoma cells to modulate the genomic distribution of BRD4 and MITF, and that BRD4 binds to MITF *in-vitro* and in cells. Mechanistically, our data suggests that the interaction between BRD4 and MITF is SETD6-mediated and BRD4 methylation-dependent and requires acetylation of MITF. We propose that the formation of this complex at chromatin has a role in transcription regulation of genes that are important for melanoma initiation and progression.

## Materials and methods

### Plasmids

All BRD4 plasmids were previously described (28). pcDNA-FLAG BRD4-N140A and pcDNA-FLAG BRD4-N140F were generated using site directed mutagenesis. Primers sequences are shown in supplementary Table 1. SETD6 plasmids were also produced in previous papers (19,21–23,26–30). Human MITF-A coding sequence was excised from plasmid pEGFP-N1-MITF-A (Addgene #38132). Human MITF-M coding sequence was excised from plasmid pCMV-Tag4A-MITF-M, which was kindly provided by the laboratory of Prof. Victoria P. Belancio. MITF coding sequences were amplified by using PCR with compatible primers as indicated in supplementary Table 1. Amplified PCR products were digested with AscI and PacI restriction enzymes and subcloned into pcDNA3.1 3×Flag and pcDNA3.1 3×HA plasmids, as well as pET-Duet-His plasmid for protein purification. KAPA HiFi HotStrart ReadyMix (KAPA Biosystems) was used for the PCR reactions. All cloned plasmids were confirmed by sequencing.

### Cell lines treatment

Melanoma cell lines SKmel147 (NRAS mutant), and Human embryonic kidney cells (HEK293T) were maintained in Dulbecco’s modified Eagle’s medium (Sigma, D5671) with 10% fetal bovine serum (FBS) (Gibco, 10270106), 1% penicillin-streptomycin (Sigma, P0781), 2 mg/ml L-glutamine (Sigma, G7513) and non-essential amino acids (Sigma, M7145). Cells were cultured at 37°C in a humidified incubator with 5% CO2.

### Transfection and Infection

Transfections of HEK293T were performed using polyethyleneimine (PEI) reagent (polyethyleneimine Inc., 23966) and Mirus reagent. TransIT-X2 (MC-MIR-6000-1.5) was used for melanoma cells (SKmel147), according to the manufacturer’s instructions.

For stable transfection in SKmel147 cell lines, retroviruses were produced by transfecting HEK293T cells with the indicated pWZL constructs (empty, Flag BRD4 wild-type 1-477aa or Flag BRD4 K99R 1-477aa) and with plasmids encoding VSV and gag-pol. Target cells were infected with the viral supernatants and selected with 250 μg/ml hygromycin B (TOKU-E), respectively.

### SETD6 knock-out by CRISPR/CAS9

For SKmel147 CRISPR/Cas9 SETD6 knock-out cells, four different gRNAs for SETD6 (supplementary Table 1) were cloned into lentiCRISPR plasmid (Addgene, #49535). 3×10^5^ melanoma cells per 6 well-plate were plated and transfected with Mirus reagent, TransIT-X2, according to the manufacturer’s protocol. Following transfection and puromycin selection (2.5μg/mL), single clones were isolated, expanded, and validated by sequencing.

### RNA-Sequencing analysis

Total RNA was extracted from SKmel147 cells (SETD6 control vs. KO) using the NucleoSpin RNA Kit (Macherey-Nagel). Samples were prepared in two biological replicates for control cells and four biological replicates for SETD6 KO cells. Libraries were prepared using the INCPM-mRNA-seq protocol. Briefly, the polyA fraction (mRNA) was purified from 500 ng of total input RNA followed by fragmentation and the generation of double-stranded cDNA. Afterward, an Agencourt Ampure XP beads cleanup (Beckman Coulter), end repair, A base addition, adapter ligation, and PCR amplification steps were performed. Libraries were quantified by Qubit (Thermo fisher scientific) and TapeStation (Agilent). Sequencing was done on a Hiseq instrument (Illumina) using two lanes of an SR60_V4 kit, allocating 20M reads per sample (single read sequencing).

### Data Processing of SETD6 KO RNA-Seq

The analysis of the raw sequence reads was carried out using the NeatSeq-Flow platform [https://doi.org/10.1101/173005]. The sequences were quality trimmed and filtered using Trim Galore (version 0.4.5) (quality cutoff=25, length cutoff=25) and cutadapt (version 1.15) [DOI: https://doi.org/10.14806/ej.17.1.200]. Alignment of the reads to the human genome (GRCh38) was done with RSEM (version 1.2.28) (31) (option ‘bowtie2’) and calculation of number of reads per gene per sample was also done with RSEM. Quality assessment of the process was carried out using FASTQC (version 0.11.8) and MultiQC (version 1.0.dev0) (32). Genes with low expression values (mean count < 1 over all samples) were excluded from differential expression analysis.

Read counts for differential gene expression were analyzed with the DESeq2 R package(33) using the the NeatSeq-Flow platform DESeq2 module. For the 4 CRISPR-treated KO samples vs. the two control samples, genes with fold change ≥ 1 and adjusted P < .05 were considered as significantly differentially expressed genes. Significant genes were clustered using the ‘eclust’ function from the factorextra R package (10.32614/CRAN.package.factoextra) with the default restriction of maximum clusters. Enrichment for Gene Ontology biological processes and KEGG pathways was performed using clusterProfiler v4.0 R package [DOI:10.1016/j.xinn.2021.100141]. RNA seq data was deposited into the Gene Expression Omnibus database under accession number GSE298160.

### ChIP Enrichment Analysis (ChEA)

ChIP-X Enrichment Analysis is a gene-set enrichment analysis tool tailored to test if query gene-sets are enriched with genes that are putative targets of transcription factors. ChEA utilizes a gene-set library with transcription factors labeling sets of putative target genes curated from published ChIP-chip, ChIP-seq, ChIP-PET, and DamID experiments (34). For SKmel147 SETD6 RNA-seq, differentially expressed genes (p < 0.05) were analyzed in the Enrichr platform for gene ontology (GO) biological processes and Hallmark gene sets.

### Colony-formation (Clonogenic) assay

Cells were seeded in six-well plates at 1000, 2000, or 5000 cells per well and incubated at 37°C incubator for 5–10 days. Cell colonies were washed twice with Phosphate buffered saline (PBS), fixed, and stained with 0.5% crystal violet in 20% methanol for 5min. Representative wells were photographed. Each experiment was performed in duplicate (technical replicates). Crystal violet staining was solubilized in 2% SDS and quantified at 550 nm using a Tecan Infinite M200 plate reader.

### Adhesion Assay

For cell adhesion assay, 3×10^5^ cells/well were plated on 96-well plate for 1 hour, followed by a PBS wash and crystal violet staining (0.5% crystal violet in 20% methanol). The crystal violet stained cells were solubilized in 2% SDS and quantified at 550 nm using a Tecan Infinite M200 plate reader.

### Antibodies and Western blot Analysis

Primary antibodies used were anti-Flag (Sigma, F1804), anti-HA (Millipore, 05–904), anti-Actin (Abcam, ab3280), anti-SETD6 (Genetex, GTX629891), anti-pan-methyl (Cell signaling, 14679), anti-pan-acetyl lysine (Cell signaling, 9681s) anti BRD4 (Bethyl, A700-004), anti BRD4 K99me1 antibody (U292-FT) (28), anti MITF (Santa Cruz, sc-515925), anti-histone 3 (H3) (Abcam, ab10799), and GFP (Abcam, ab290). HRP-conjugated secondary antibodies, goat anti-rabbit and goat anti-mouse, were purchased from Jackson ImmunoResearch (111-035-144, 115-035-062, respectively). For Western blot analysis, cells were homogenized and lysed in radioimmunoprecipitation assay (RIPA) buffer [50 mM tris-HCl (pH 8), 150 mM NaCl, 1% Nonidet P-40, 0.5% sodium deoxycholate, 0.1% SDS, 1 mM dithiothreitol (DTT), and 1:100 protease inhibitor mixture (Sigma)], for 10 min on ice. Proteins denatured with Laemmli sample buffer (250mM Tris-HCl pH 6.8, 10% SDS, 30% glycerol, 5% β-mercaptoethanol, a pinch of bromophenol blue), boiled for 5min at 95°C and were resolved on 8%-12% SDS–polyacrylamide gel electrophoresis (PAGE), followed by blotting onto PVDF membranes and analysis with appropriate antibodies.

### Chromatin Extraction by Biochemical Fractionation and protein-protein interaction analysis by immunoprecipitations (IP)

Cells were cross-linked using 1% formaldehyde (Sigma) added directly to the medium and incubated on a shaking platform for 10min at room temperature. The cross-linking reaction was stopped by adding glycine to a final concentration of 0.125M and incubating for an additional 5min on the shaking platform. Cells were washed twice with PBS and then lysed in 1 ml cell lysis buffer (20 mM Tris-HCl pH 8, 85 mM KCl, 0.5% Nonidet P-40, 1:100 protease inhibitor cocktail) for 10 min on ice. Nuclear pellets were resuspended in 120-200μl nuclei lysis buffer (50mM Tris-HCl pH=8, 10mM EDTA, 1% SDS, 1:100 protease inhibitor cocktail) for 10min on ice, and then sonicated (Bioruptor, Diagenode) at high power settings for 3 cycles, 6min each (30sec ON/OFF) or sonicated by focused-ultrasonicator (ME220, Covaris). Samples were centrifuged (20,000g, 15 min, 4°C) and the soluble chromatin fraction was collected. For protein-protein interaction analysis with endogenous BRD4 or MITF, the soluble chromatin was pre-cleared with Magna ChIP™ Protein A+G Magnetic Beads (Millipore, 16-663) for 1h. Then 3 or 6μL of Anti-BRD4/MITF antibody was added to the pre-cleared sample for each IP reaction and the tubes were rotated in a shaker at 4°C overnight. Then, 20μL Magna ChIP™ Protein A+G Magnetic Beads were added to each IP reaction, followed by incubation in a shaker at 4 °C for 2h. The tubes were then placed on a magnetic separator to capture magnetic beads and the associated proteins. The supernatants were removed, and the magnetic beads were washed with 1mL of PBSx1. For pan-methyl immunoprecipitation, cell lysates in RIPA buffer were pre-cleared with A/G agarose beads (Santa Cruz, SC-2003) for 1h and then incubated overnight at 4°C with pan-methyl antibody which was pre-conjugated to A/G agarose beads. The immunoprecipitated complexes were washed once with PBS and then resolved in protein sample buffer and analyzed by Western blot. For SAHA treatment, cells were treated for 4h with 5-20μM compound or with (add percentage) dimethyl sulfoxide as a vehicle control. SAHA was provided by Dr. D. Toiber (Ben-Gurion University, Israel). For JQ1 treatment, lysates of cells treated with 1μM JQ1 and incubated with primary antibody for IP. JQ1 was provided by Dr. V. Shoshan-Barmatz (Ben-Gurion University, Israel).

### Recombinant protein purification

*Escherichia coli* BL21 transformed with a plasmid expressing a His-tagged protein of interest was grown in LB media. Bacteria were harvested by centrifugation after IPTG induction and lysed by sonication on ice (25% amplitude, 1 min total, 10/5 sec ON/OFF). Homogenized with ice-cold lysis buffer containing phosphate-buffered saline (PBS), 10mM imidazole, 0.1% Triton X-100, 1mM PMSF, and one complete mini protease inhibitors tablet (Roche). After adding 0.25mg/ml lysozyme for 30 min, the lysates were subjected to sonication on ice (18% amplitude, 1 min total, and 10 sec ON/OFF). The tagged fusion proteins were purified on a His-Trap column using an AKTA Pure protein purification system (GE). The proteins were eluted with 0.5M imidazole in PBS buffer, followed by overnight dialysis (PBS, 10% glycerol).

### In vitro methylation assay

The methylation assay reaction (total volume of 25μl) contained 1μg of His-Sumo BRD4, 4μg His-SETD6, 1μl of 2mCi of 3H-labeled S-adenosyl-methionine (SAM) (Perkin-Elmer, AdoMet) and PKMT buffer (20mM Tris-HCl pH 8, 10% glycerol, 20mM KCl, 5mM MgCl_2_). The reaction tubes were incubated overnight at 30°C. Then, the reactions were resolved by SDS-PAGE for Coomassie staining (Expedeon, InstantBlue) or autoradiography (Typhoon FLA 7000, GE).

### Enzyme-linked immunosorbent assay (ELISA)

2μg of recombinant proteins (BSA, MBP-RelA, His-Sumo BRD4, and His MITF) in PBS were added to a sticky surface (Greiner Microlon) 96-well plate and incubated for 1h at room temperature (RT) followed by 3% BSA blocking in 1X Phosphate-Buffered Saline, 0.1% Tween (PBST) for over-night incubation at RT. After 3 washes with PBST, 0.5μg GST-SETD6 or GST protein (negative control) diluted in 1% BSA in PBST was added to the wells for 1h at RT. Plates were washed and incubated with primary antibody (anti-GST, 1:4000 dilution) followed by incubation with an HRP-conjugated secondary antibody (goat anti-rabbit, 1:2000 dilution) for 1 hr. Finally, TMB reagent and then 1N H_2_SO_4_ were added; the absorbance at 450nm was detected using a Tecan Infinite M200 plate reader. Results are represented as relative absorbance compared to GST or BSA\ His-Sumo.

### Cleavage under targets & release using nuclease (CUT&RUN)

Cut&Run was performed using the CUTANA™ Assay kit (EpiCypher) according to the manufacturer’s instructions. An input of 1×10^6^ cells per sample was processed according to the manufacturer’s protocol. More specifically, with the use of nuclear extraction buffer the Cut&Run assay was performed directly on nuclei. Digitonin was used at a final concentration of 0.01% for nuclear permeabilization. 0.5μg of controls antibodies (IgG and H3K4me3) were used as recommended and 1μg of anti-BRD4 (EpiCypher 13-2003), anti-MITF (Santa Cruz, sc-515925) and anti-Flag (Sigma, F1804) antibodies were used per sample. Libraries were quantified with the Qubit dsDNA HS Assay Kit and DNA fragment sizes were assessed using an Agilent TapeStation. Resulting libraries were sequenced with 50-150bp paired-end reads, with 15M reads per sample using a NextSeq 2000 system from Illumina through the Center for Advance Genomics, IKI, BGU.

### CUT&RUN data analysis

The analysis was carried out using the NeatSeq-Flow platform (35) [https://doi.org/10.1101/173005]. Raw sequencing data in fastq format was trimmed by Trim Galore v0.4.5 (length= 25, q-25), reads that were too short were discarded. FastQC v0.12.0 was used for reads quality control. BWA Mapper (version 0.7.12, default parameters t=20, B=5) (36) was used to align paired-end reads to reference human genome assembly hg38 and to the spike-in control (*Escherichia coli*) reference genome assembly. SAMtools (37) was used to remove PCR duplicates and create coordinate-sorted BAM files. Normalization between samples and IgG was performed with a custom script that calculates read scaling factor based on spike-in DNA reads of each sample. The scaling factor was then used in the bamCoverage v3.5.6 by deepTools (38) to normalize each sample according to its spike-in, Bigwig files were created by bamCoverage and reads mapped to blacklisted areas designated by ENCODE (39) were filtered. Reads that refer to off-chromosome locations were removed with BedClip v377 and sorted by Bedtools v2.30.0 (40). Bigwig files were then created with bedGraphToBigWig by deepTools (38). For BRD4, peaks were called by MACS (version 3.0.0) (41) (“callpeak” function, options “nomodel”, “broadpeak”, broad-cutoff = 0.05). For MITF, peaks were called by SEACR v1.3 (using the top 5 percentile as the high-confidence peak calibration in “stringent” mode) (42). Peaks within 100 bp were then merged and sorted with Bedtools.

The union of peaks from each two replicates (BRD4, Flag-BRD4 and MITF) were then intersected by Bedtools (version 2.31.0)(40), resulting in a peak list that included only peaks that exist in both replicates. Peaks were viewed in IGV (version 2.16.1)(43) and annotated with ChIPseeker (44,45). Matrices were generated with DeepTools createMatrix, and Heatmaps were plotted with deepTools plotHeatmap (38). Cut&Run data was deposited into the Gene Expression Omnibus database under accession number GSE298161.

### Proximity Ligation Assay (PLA)

Cells were cultivated on coated coverslips, washed with PBS and fixed in cold 4% Paraformaldehyde (PFA) at RT for 15min. Cell permeabilization was performed using 0.5% Triton X-100 in PBS for 10min in RT. PLA (Duolink) was performed according to the manufacturer’s instructions (Sigma) using antibodies against BRD4 (Bethyl, A700-004), MITF (Santa Cruz, sc-515925) and Flag (Sigma, F1804) overnight in 4°C. Images were acquired by confocal Spinning disk microscopy with a 100x or 63x objective. Each frame represents maximum intensity projection for Z-stacks captured, and 4-6 frames were captured for each sample. The PLA units were calculated per cell as the ratio between the number of dots within the nucleus and the nucleus area (stained with DAPI, using Duolink mounting media). Each nucleus is then represented as a point in the quantification graph.

### PLA image processing and data analysis

For the automated counting of PLA emitters per nucleus in batch mode, an ImageJ (Fiji) macro was implemented (46). In each image, the nuclei were first segmented by applying a Gaussian blur and using standard thresholding algorithms, such as Otsu. The segmented nuclei were then converted into ImageJ ROIs using the MorpholibJ (47) and PTBIOP [https://www.epfl.ch/research/facilities/ptbiop/image-processing/] plugins. For each ROI (nucleus), the number of PLA emitters was determined by identifying local maxima following blob detection via the Difference of Gaussian (DoG) algorithm.

### Quantification and Statistical Analyses

Statistical analyses for all assays were performed with GraphPad Prism software, using one-way, two-way analysis of variance (ANOVA) or student’s t-test.

## Results

### SETD6 Modulates Gene Expression and Cellular properties in Melanoma Cells

Kaplan-Meier survival analysis of 367 melanoma patients with low (n=282) or high (n=85) expression of SETD6 revealed that a high level of SETD6 correlates with poor patient prognosis **(Fig. 1A)**. These findings suggest that SETD6 may have a functional role in melanoma pathobiology. To test SETD6’s role in transcription regulation in melanoma, we performed RNA-sequencing experiments for control (CT) and knock-out (KO) of SETD6 in SKmel147 melanoma cells (4 independent gRNA clones), which we have generated and validated by sequencing **(Fig. S1)**. 52 down-regulated genes and 169 up-regulated genes were observed upon depletion of SETD6 **(Fig. 1B)**. Gene Ontology (GO) analysis revealed that the differential expressed genes (DEGs) are enriched in pathways such as cell proliferation and cell adhesion, two processes which are linked to melanoma as well as SETD6 enzymatic activity (48–50) **(Fig. 1C)**. Consistent with these observations, knockout of SETD6 in SKmel147 cells attenuates colony formation compared to the control cells **(Fig. 1D)**. In a cell adhesion assay, we observed a significant decrease in the adhesive properties of SETD6-depleted cells compared to the control cells **(Fig. 1E)**. Taken together, these results indicate that SETD6 influences gene expression profiles in melanoma cells, thereby promoting colony formation and cell adhesion.

**Figure 1.**
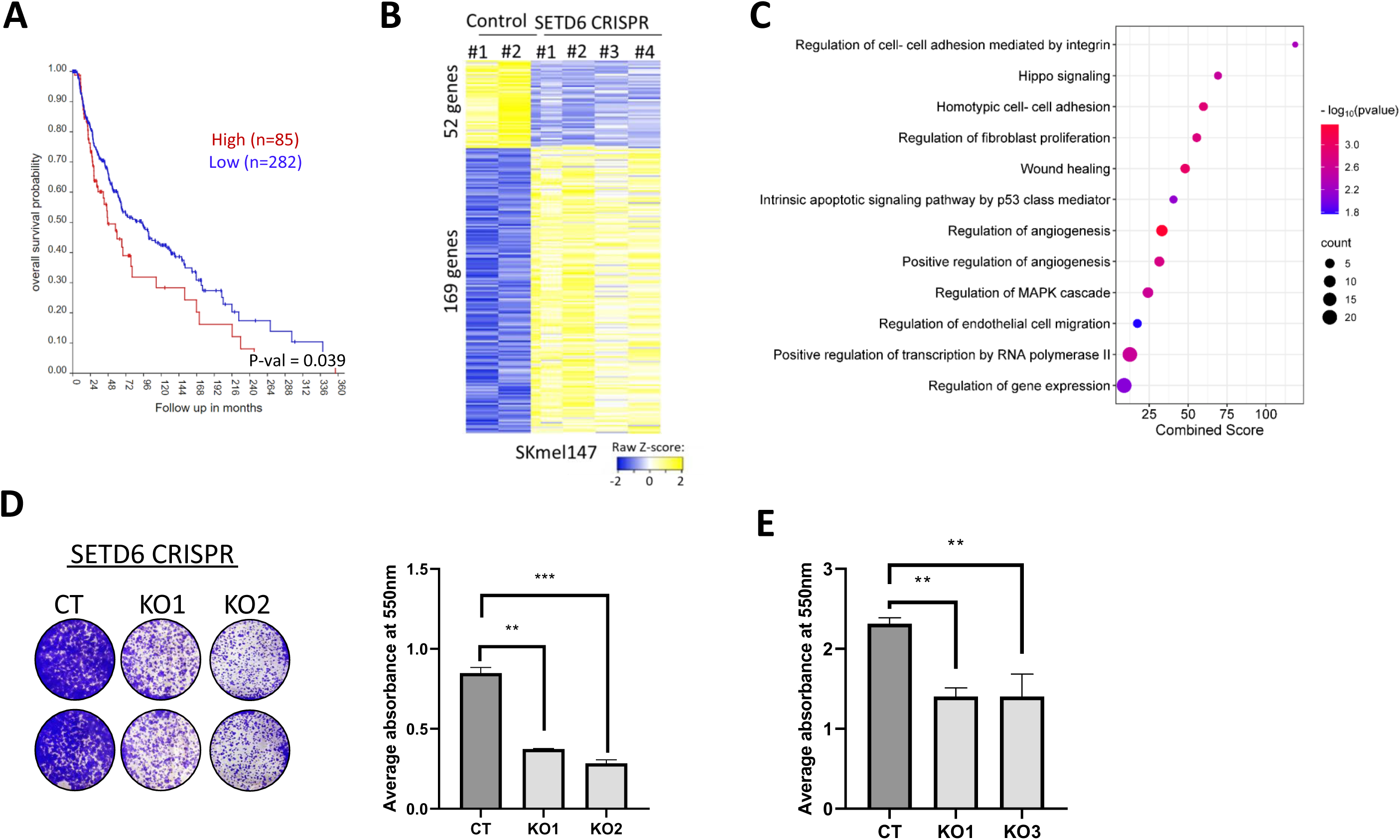
SETD6 regulates gene expression and cellular properties of melanoma cells. **(A)** Kaplan-Meier survival curve for high vs. low levels of SETD6 in melanoma patients (TCGA database, p=0.039), created using R2 genomics analysis and visualization platform [https://hgserver1.amc.nl/]. **(B)** Heatmap showing up- and down-regulated genes from RNA-sequencing analysis of 2 SETD6 control and 4 KO SKmel147 cells independent clones. Yellow and blue colors represent high and low expression levels, respectively. **(C)** Selected pathways from gene ontology (GO) analysis of the differentially expressed genes were analyzed. Circle size represents the count of differentially expressed genes related to each pathway. **(D)** Colony-formation assay for SETD6 CT and KO SKmel147 cells. Images of colonies stained with Crystal violet; and Crystal violet-stained cells were dissolved in 2% SDS and the absorbance at 550nm was measured, whereby error bars represent the SEM. Statistical analysis was performed for two experimental repeats using one-way ANOVA (** p<0.01). **(E)** Adhesion assay for SKmel147 parental, SETD6 CRISPR CT and KO cells. Adherent cells stained with Crystal violet, dissolved and analyzed as described in D.

### SETD6 interacts with and methylates BRD4 in melanoma cells

In a recent study from our lab, we found that SETD6 methylates chromatin-bound BRD4 on lysine-99 (K99). This methylation influences gene expression by selectively recruiting the transcription factor E2F1 to its target genes in breast cancer cells (28). Kaplan-Meier survival analysis of 367 melanoma patients with low (n=342) or high (n=25) expression of BRD4 revealed that a high level of BRD4 expression is correlated with poor patient prognosis **(Fig. 2A)**. Interestingly, this is in agreement with the same trend observed for SETD6 (**Fig. 1A**). These findings raised the possibility that SETD6 methylation of BRD4 at K99 might occur in melanoma cells. To address this hypothesis, we first immunoprecipitated (IP) endogenous BRD4 in melanoma cells and confirmed its interaction with endogenous SETD6 at chromatin **(Fig. 2B)**. Next, to test if BRD4 is a substrate for methylation by SETD6 in melanoma, we used a lysine pan-methyl antibody in control and SETD6 KO cells. As shown in **Figure 2C**, our results suggest that over-expressed Flag-BRD4 is methylated in a SETD6-dependent manner. To test if BRD4 is methylated on K99, we probed cell lysate derived from SETD6 control and KOs cells with BRD4 K99me1 antibody (28) **(Fig. 2D).** We observed that the level of BRD4K99me1 was reduced in the SETD6 knockout cells. The residual signal observed in the SETD6 depleted cells (**Fig. 2D**, lanes 2 and 3) comes from the endogenous BRD4 protein which is still present. To validate this observation, we utilized a lysine pan-methyl antibody in both Flag-BRD4 WT and Flag-BRD4 K99R mutant overexpressed cells. The results revealed a reduction in the methylation of BRD4 K99R mutant compared to WT **(Fig. S2A)**. Phenotypically, colony formation assay was performed to test melanoma cell proliferation using SKmel147 stably expressing either empty vector, BRD4 WT or BRD4 K99R mutant which cannot be methylated by SETD6 **(Fig. 2E and Fig. S2B** for expression validation). SKmel147 cells expressing BRD4 WT demonstrate augmented colony formation compared to the empty vector control. Consistent with the experiments shown in **Figure 1D** where SETD6 was depleted, cells expressing BRD4 K99R formed fewer colonies compared to BRD4 WT cells. We also observed a significant decrease in the adhesion ability of BRD4 K99R cells compared to the BRD4 WT-expressing cells **(Fig. 2F)**. Altogether, these findings suggest that SETD6 binds and methylates BRD4 at K99 in melanoma cells to positively regulate cell proliferation and adhesion.

**Figure 2.**
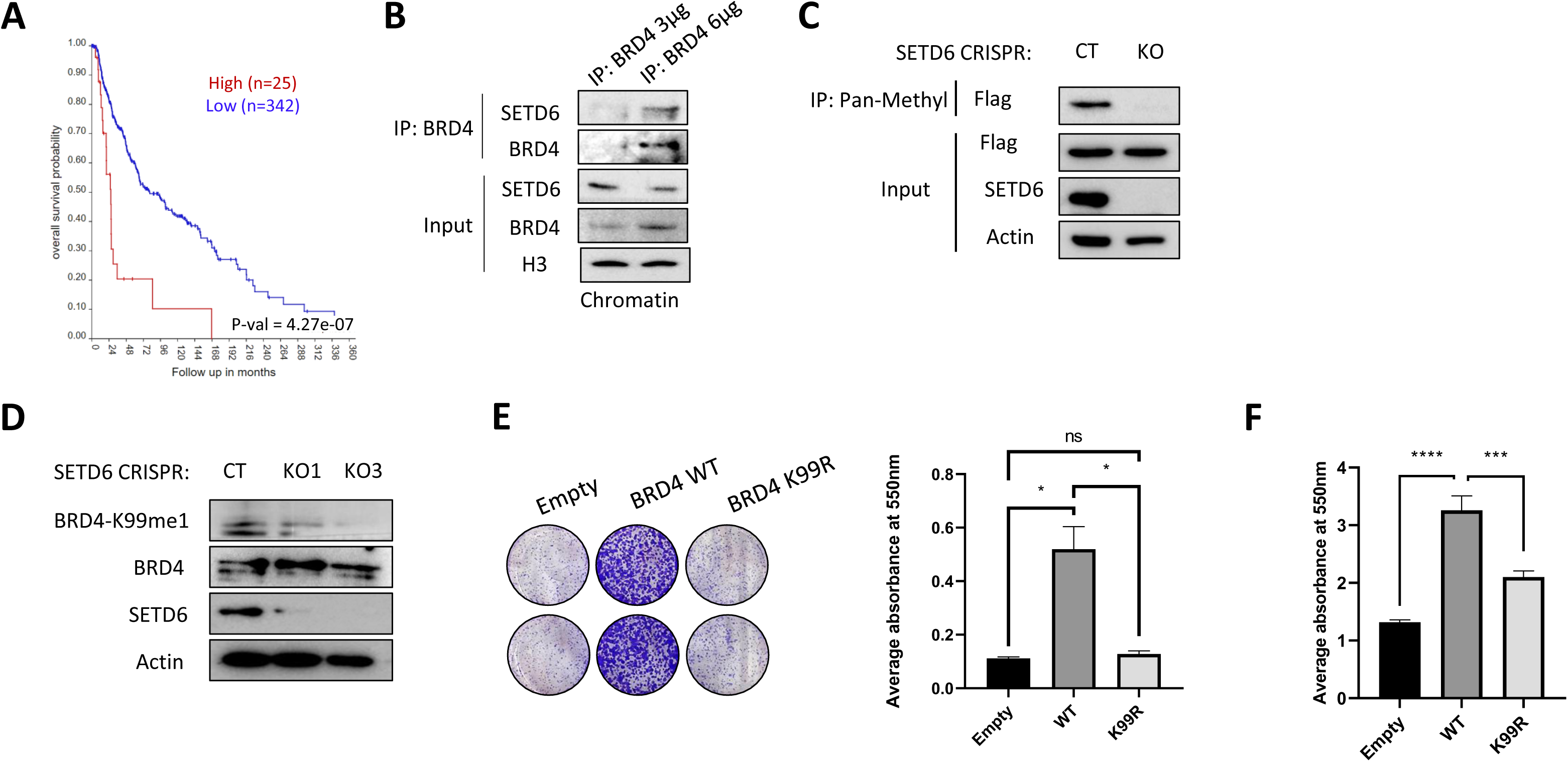
SETD6 interacts with BRD4 and methylate it in melanoma cells. **(A)** Kaplan-Meier survival curve for high vs. low levels of BRD4 in melanoma patients (TCGA database, p=4.27e-07) created using R2 genomics analysis and visualization platform. **(B)** Chromatin extract from SKmel147 cells were immunoprecipitated with anti-BRD4 antibody followed by WB with the indicated antibodies. Input - levels of BRD4, SETD6, and Histone-3 (loading control) in the total chromatin extracts. **(C)** Cellular methylation assay wherebySETD6 CT and KO cells were transfected with Flag BRD4 WT or Flag BRD4 K99R plasmids. Cell lysates were immunoprecipitated with pre-conjugated pan-methyl A/G agarose beads, and proteins in the immunoprecipitateand input samples were detected by Western blot with indicated antibodies. **(D)** WB analysis for CT and two SETD6 KO cells (KO1 and KO3) with the indicated antibodies. BRD4-K99me1 (U292-FT) – an antibody which specifically recognizes methylated BRD4 at K99. **(E)** Colony-formation assay for SKmel147 cells stably expressing Empty, Flag-BRD4 WT and Flag-BRD4 K99R. Error bars represent the SEM. Statistical analysis was performed for two experimental repeats using one-way ANOVA (ns = not significant, *p<0.05). **(F)** Adhesion assay for SKmel147 cells stably expressing Empty, Flag-BRD4 WT and Flag-BRD4 K99R. Adherent cells stained with Crystal violet, dissolved and analyzed as described in F (*** p<0.001, **** p<0.0001).

### BRD4 K99 methylation by SETD6 regulates its genomic occupancy

Given that BRD4 is a transcription regulator (6), we examined whether SETD6 regulates endogenous BRD4 genomic occupancy. To do so, we performed Cleavage Under Targets and Release Using Nuclease (CUT&RUN) assays (51) in SKmel147 cells control and SETD6 KO cells **(Fig. 3A)**. We found a significant decrease of BRD4 enrichment in SETD6 KO cells which may suggest that this effect is mediated by BRD4 methylation at K99. To test this hypothesis, we performed a CUT&RUN using Flag antibody for cells stably expressing Flag-BRD4 WT or K99R mutant **(Fig. 3B)**. Consistently, we found a significant decrease of enrichment in BRD4 K99R cells compared to BRD4 WT. To determine the genomic locations which are SETD6 and BRD4 K99me dependent, we intersected the annotated peaks of endogenous BRD4 in SETD6 control cells (Fig. 3A) with overexpressed Flag-BRD4 WT cells (Fig. 3B). The analysis revealed 364 significant shared loci **(Fig. 3C)**. Genome browser view of the CUT&RUN experiments is presented in **Figure 3D**. ChIP-seq tracks for BRD4 (GSM2476358) were added for comparison. These results demonstrate that BRD4 methylation at K99 by SETD6 affects its genomic distribution and is linked to the regulation of oncogenic related pathways.

**Figure 3.**
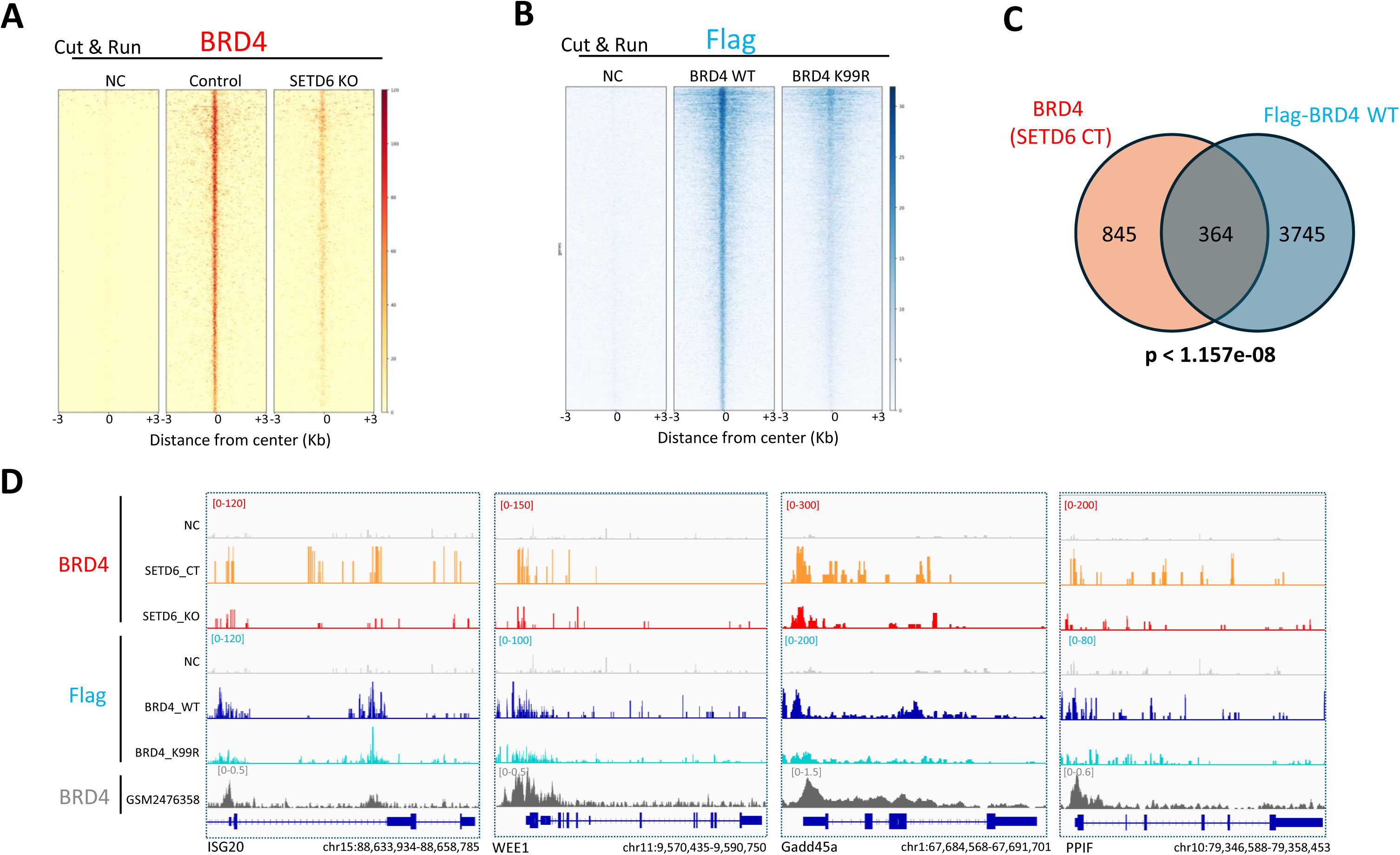
BRD4 K99-methylation by SETD6 affects its genomic distribution. **(A)** Heatmaps showing CUT&RUN read densities for IgG (NC) and BRD4 across BRD4 genomic peaks in SETD6 control and KO SKmel147 cells. **(B)** Heatmaps showing CUT&RUN read densities for IgG (NC) and Flag-BRD4 across Flag-BRD4 genomic peaks in SKmel147 cells expressing Flag-BRD4 WT or Flag-BRD4 K99R. **(C)** Venn diagram showing common genes for BRD4 in SETD6 CT cells and Flag in BRD4 WT cells as identified in the CUT&RUN analyses. **(D)** Representative binding profiles of four genomic regions.

### SETD6 regulates MITF genomic distribution

ChIP enrichment analysis (ChEA) (52) analysis on the 364 shared loci (Fig. 3C) revealed significant enrichment of several transcription factor **(Fig. 4A)**. As expected BRD4 was among the 4 most significant ones. Both NFKB1 and NR3C1 were shown before to be associated with BRD4 cellular activity (53–55). We were particularly interested in MITF which is a crucial regulator of melanocyte development, controlling differentiation, cell-cycle progression, pigmentation, and melanocyte survival (1). It is also identified as an amplified oncogene in some human melanomas (1). Interestingly, a direct interaction between recombinant SETD6 and MITF was observed using an Enzyme-Linked Immunosorbent Assay (ELISA) **(Fig. S3A)**, RelA served as positive control for the experiment (24). The physical interaction between SETD6 and MITF was also validated in cells through an IP experiment at the chromatin level **(Fig. S3B)**. These findings raised the hypothesis that SETD6 might regulate also MITF genomic distribution and activity in melanoma cells. To address this hypothesis, we performed a CUT&RUN experiment for endogenous MITF in CT and SETD6 KO cells **(Fig. 4B)**. The results suggest that the knockout of SETD6 reduces MITF recruitment or retention to its genomic binding sites. Intersection of the annotated peaks for BRD4, Flag-BRD4 and MITF revealed 200 shared genes **(Fig. 4C)**. A snapshot for several peaks is presented in **Figure 4D**. KEGG analysis for these shared genes revealed significant enrichment for genes involved in cell adhesion **(Fig. S4)**. These findings may suggest a functional interplay between BRD4, MITF and SETD6.

**Figure 4.**
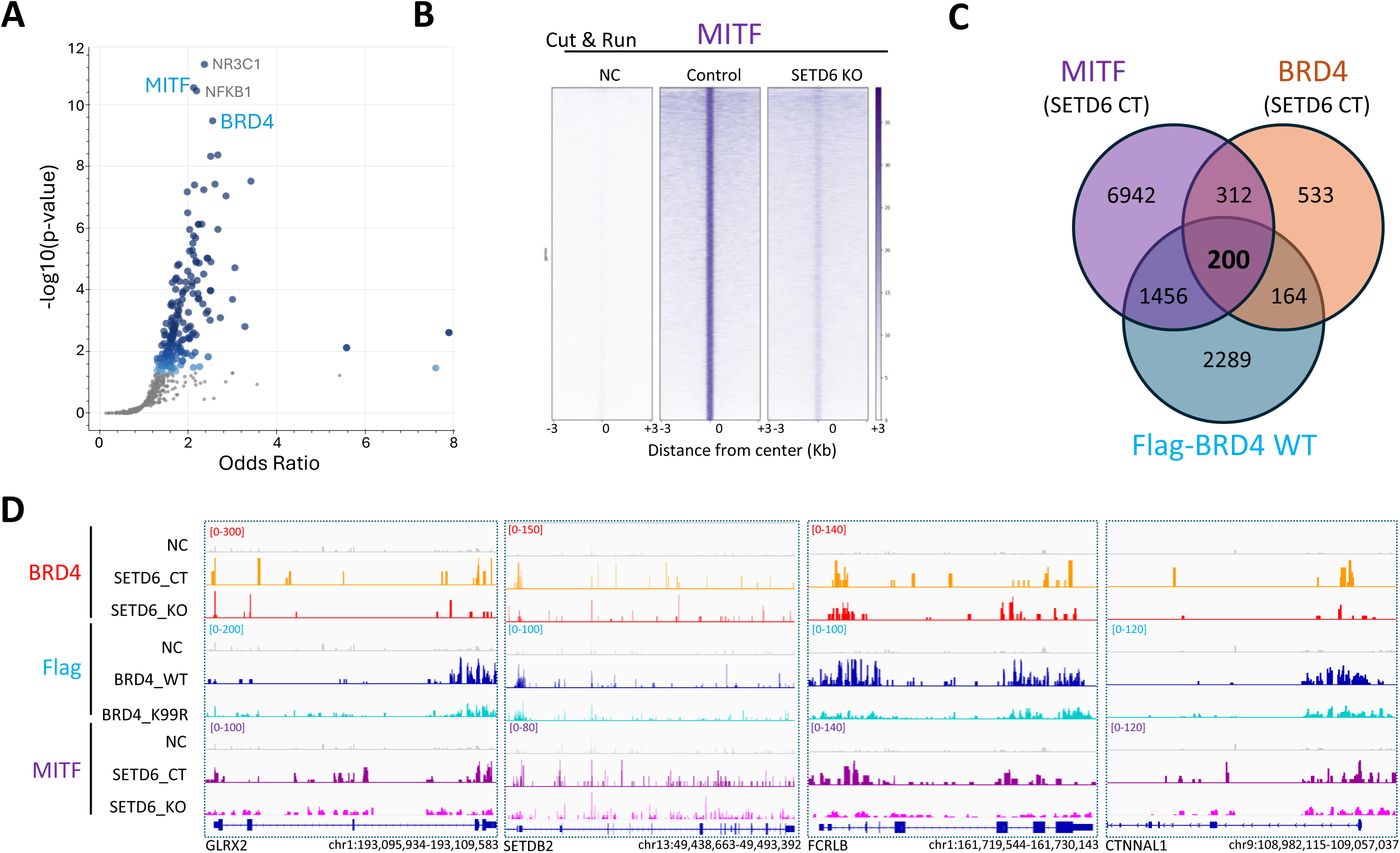
SETD6 interacts with MITF and regulates its genomic distribution. **(A)** Chip-Enrichment Analysis (ChEA) for the 364 shared peaks presented in Figure 3C. Blue points represent a significant transcription factor. Smaller gray points represent non-significant terms. **(B)** Heatmaps showing CUT&RUN read densities for IgG (NC) and MITF across MITF genomic peaks in SETD6 control and KO SKmel147 cells. **(C)** Venn diagram showing common genes for BRD4, MITF in SETD6 CT cells and Flag-BRD4 in Flag-BRD4 WT cells as identified in the CUT&RUN analysis. **(D)** Representative binding profiles of three genomic regions.

### The physical interaction between BRD4 and MITF is mediated by SETD6

To address this hypothesis, we decided to check if BRD4 interacts with MITF. We confirmed that the physical interaction between endogenous BRD4 and over-expressed Flag-MITF occurs in cells using a proximity ligation assay (PLA) **(Fig. 5A)**. Using a complementary approach, we found that endogenous MITF co-immunoprecipitated with endogenous chromatin-bound BRD4 **(Fig. S5)**. We hypothesized that the physical interaction between the proteins depends on the presence of SETD6. To address this possibility, we performed a PLA experiment using specific antibodies to endogenous BRD4 and MITF in SETD6 control and KO cells **(Fig. 5B)**. These experiments revealed that SETD6 knockout abolishes the interaction between BRD4 and MITF, suggesting that the interaction between these proteins in cells is SETD6-dependent. Given that SETD6 promotes BRD4-K99me1, we hypothesized that this specific methylation event may regulate the interaction between BRD4 and MITF. Therefore, we monitored the interaction of endogenous MITF with BRD4 WT or BRD4 K99R mutant using PLA **(Fig. 5C)**. As expected, we observed a significant reduction of PLA signal in BRD4 K99R cells compared to the BRD4 WT cells, indicating that the methylation of BRD4 at K99 positively regulates its physical interaction with MITF. To support these findings, we performed an immunoprecipitation using either BRD4 WT or K99R mutant and observed a reduced interaction between BRD4 and MITF in the mutant **(Fig. 5D)**. Taking together, these results suggest that the physical interaction between BRD4 and MITF is SETD6 and BRD4-K99 methylation dependent.

**Figure 5.**
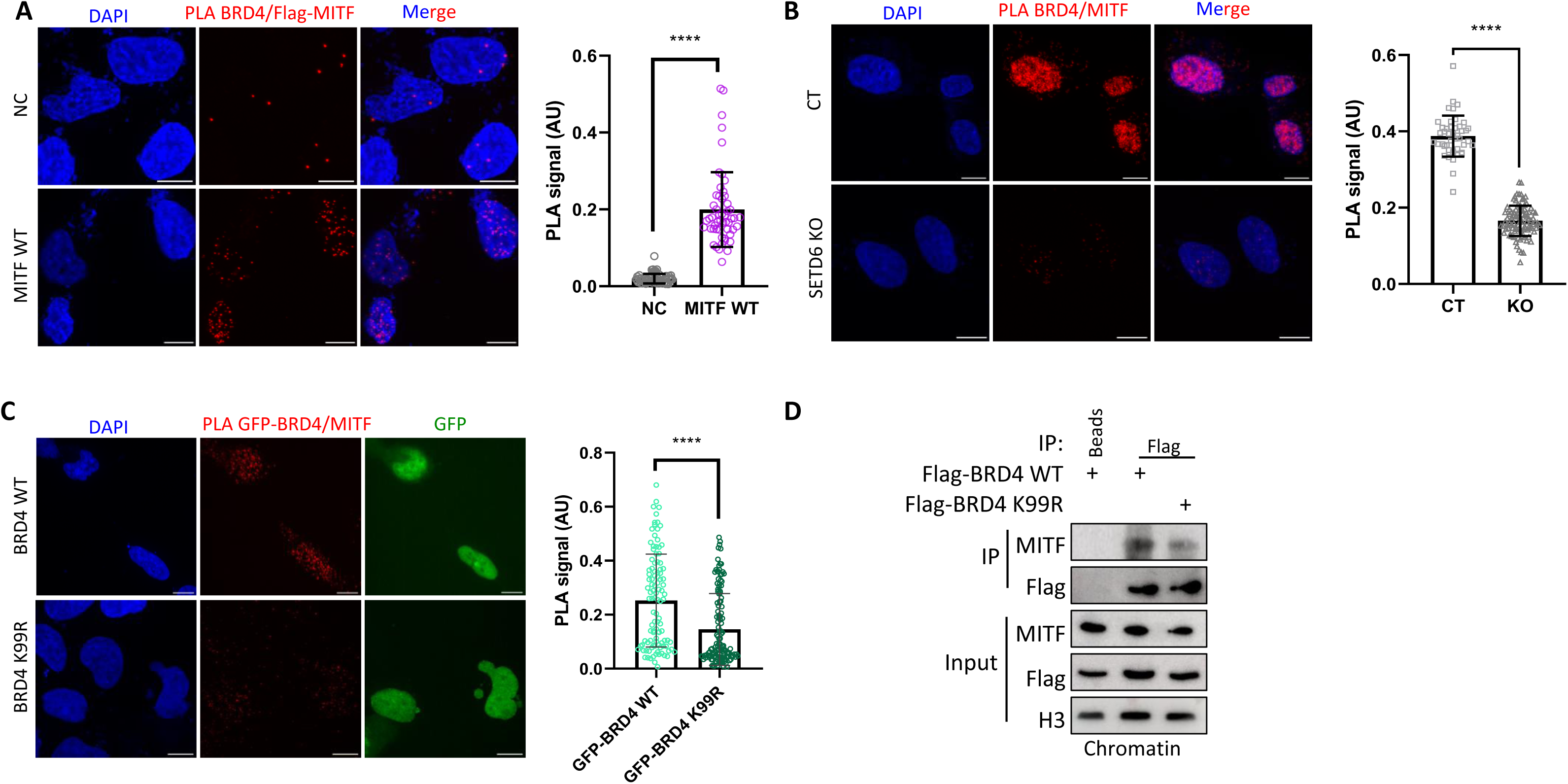
BRD4 interacts with MITF in a SETD6-dependant manner. **(A)** Left-Representative images and signal quantification (PLA dots per nucleus, AU) of PLA detecting BRD4 and Flag-MITF proximity in SKmel147 cells compared to negative control (NC) with no primary BRD4 antibody. Red dots represent PLA signal for MITF-BRD4 proximity. Scale bar = 10 micron. Right-Quantification of PLA signal per sample. Statistical analysis was performed using student’s t-test (**** p<0.0001). **(B)** Left-Representative images of PLA detecting BRD4 and MITF proximity in SKmel147 SETD6 CT and KO cells. Red dots represent PLA signal for MITF-BRD4 proximity. Scale bar = 10 micron. Right-PLA signal quantification (PLA dots per nucleus, AU) for each sample. Statistical analysis was performed using student’s t-test (**** p<0.0001). **(C)** Left-Representative images and signal quantification (PLA dots per nucleus, AU) of PLA detecting GFP-BRD4 and MITF in SKmel147 cells overexpressed GFP-BRD4 WT and GFP-BRD4 K99R. Red dots represent PLA signal for MITF-BRD4 proximity. Scale bar = 10 micron. Right-Quantification of PLA signal per sample. Statistical analysis was performed using student’s t-test (**** p<0.0001). **(D)** Chromatin extract from SKmel147 cells overexpressing Flag-BRD4 WT or K99R were immunoprecipitated with anti-Flag antibody followed by WB with the indicated antibodies. Input - levels of MITF, Flag, and Histone-3 (loading control) in the total chromatin extracts.

### The bromodomain of BRD4 binds to acetylated MITF

Since MITF is known to be acetylated on several lysine residues (56,57), we hypothesized that the BRD4-MITF interaction is mediated by the bromodomain (BD) of BRD4 (6,7) and acetylated MITF. The usage of a Pan-Acetyl antibody in IP experiments allowed us to determine that MITF is indeed acetylated in SKmel147 cells **(Fig. 6A)**. To test if the interaction with BRD4 is mediated by the acetylation of MITF, we performed a PLA experiment in the presence of increasing concentrations of SAHA, a potent inhibitor of histone deacetylases (HDACs) (58). As shown in **Figure 6B**, we observed a stronger interaction between BRD4 and MITF in a SAHA dose dependent manner. Next, we performed an IP experiment in the absence or presence of the BD inhibitor, JQ1(59), to inhibit BRD4 ability to bind its acetylated partners. The results demonstrate that the interaction between over-expressed Flag-MITF and endogenous BRD4 decreased in the presence of JQ1 **(Fig. 7A).** In a reciprocal experiment, the interaction between overexpressed GFP-BRD4 and endogenous MITF decreased in the presence of JQ1 **(Fig. S6)**. These experiments indicate that the BD of BRD4 mediates the interaction with MITF.

**Figure 6.**
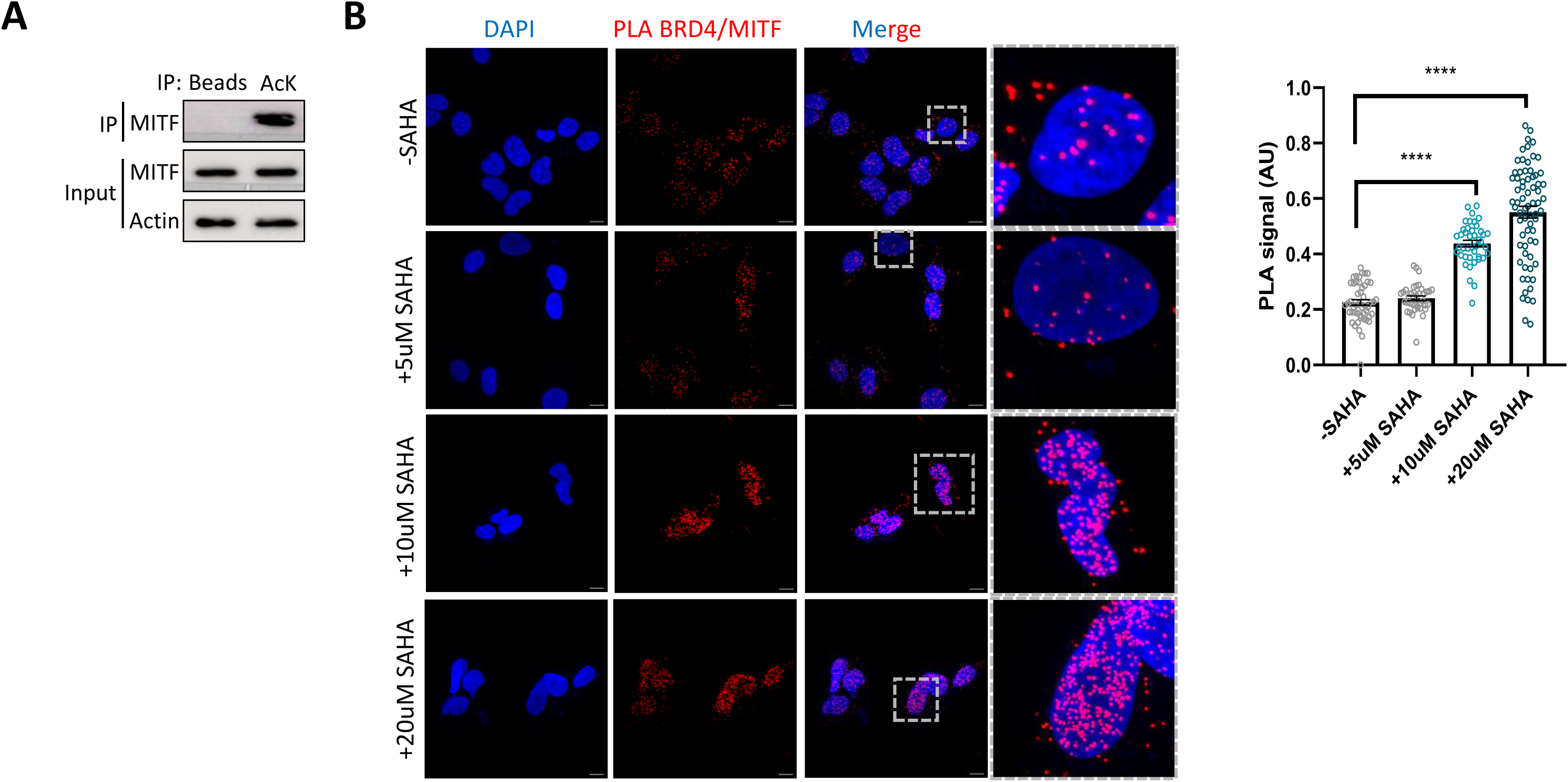
Acetylation of MITF increases its interaction with BRD4. **(A)** Acetylation assay in cells. SKmel147 cell lysates were subject to immunoprecipitation with pre-conjugated pan-acetyl lysine A/G agarose beads. Proteins in the immunoprecipitate and input samples were detected by Western blot (WB) with the indicated antibodies. **(B)** Representative images and signal quantification (PLA dots per nucleus, AU) of PLA detecting BRD4 and MITF proximity in SKmel147 cells that were untreated or treated with SAHA for 4h. Red dots represent a PLA signal for MITF-BRD4 proximity. Scale bar = 10 micron. Right-Quantification of PLA signal per sample. Statistical analysis was performed using student’s t-test (**** p<0.0001).

**Figure 7.**
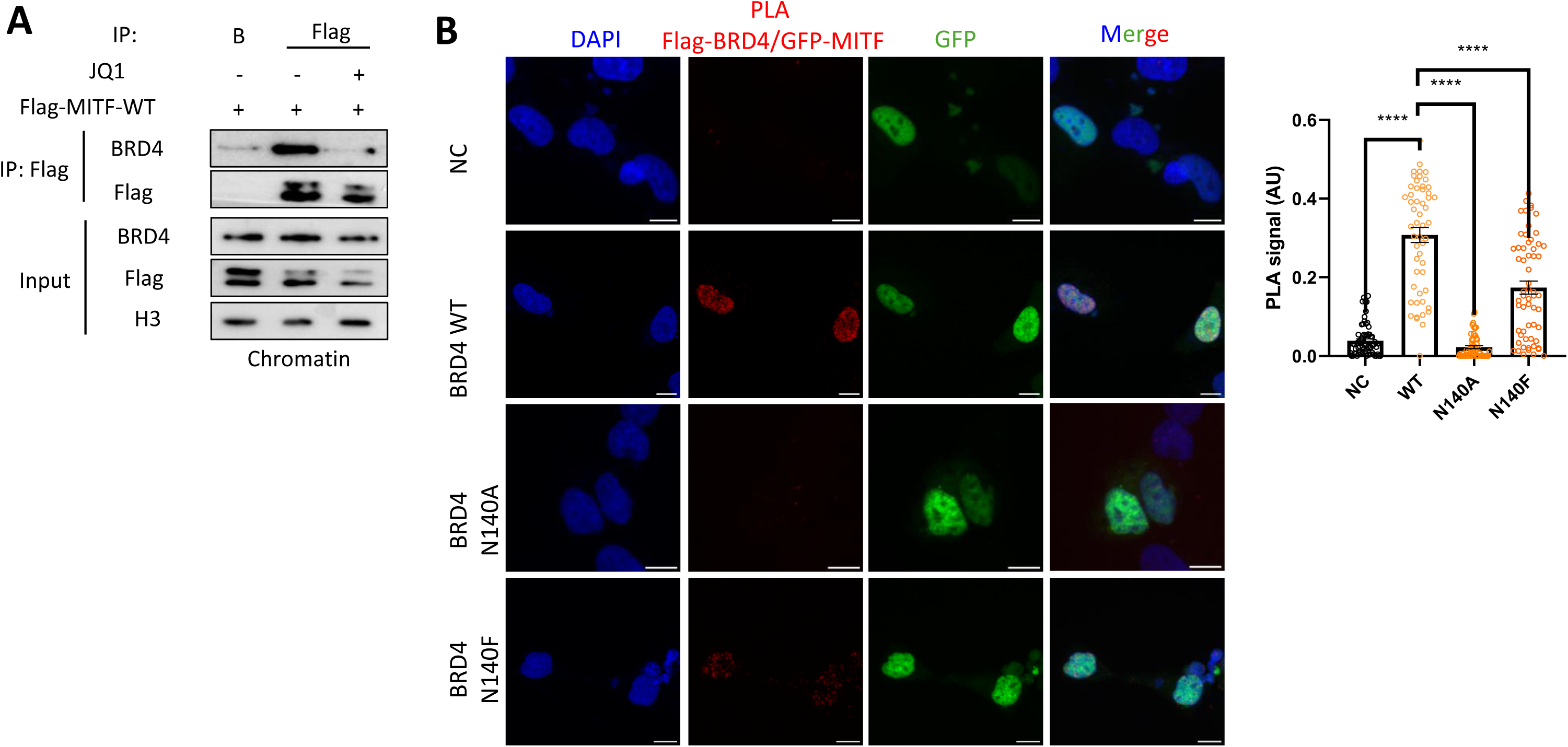
The bromodomain of BRD4 binds to acetylated MITF. **(A)** SKmel147 cells were transfected with FLAG-MITF-WT and treated with 1µM JQ1 where indicated. Chromatin fractions were then immunoprecipitated (IP) with a Flag antibody followed by WB with the indicated antibodies. Beads (B) served as negative control for the IP. **(B)** Representative images and signal quantification (PLA dots per nucleus, AU) of PLA to detect the proximity of Flag-BRD4 and MITF in SKmel147 cells overexpressing Flag-BRD4 WT, Flag-BRD4 N140A or Flag-BRD4 N140F mutants. Red dots represent a PLA signal for MITF-BRD4 proximity. Scale bar = 10 micron. Right-Quantification of PLA signal per sample. Statistical analysis was performed using student’s t-test (**** p<0.0001).

The N140 residue of BRD4 is located within its first bromodomain, and mutations at this position have previously been shown to impair BRD4 function. Notably, the well-characterized inhibitor JQ1 has been reported to form a direct hydrogen bond with N140, highlighting its importance in ligand binding. Both N140A and N140F mutants seem to reduce BRD4 affinity for acetylated lysine (60). To test the effect of these mutants on BRD4 ability to interact with MITF, we performed PLA experiment **(Fig. 7B)**. We observed a significant reduction of PLA signal in both BRD4 N140A and N140F cells compared to the BRD4 WT cells, which provides further evidence that the interaction between MITF and BRD4 occurs through the BRD4 bromodomain. Taken together, our findings suggest a model by which the interaction between MITF and BRD4 is mediated by the acetylation of MITF and the BD of BRD4 in a SETD6 dependent manner **(Fig. 8)**. We suggest that the formation of this complex at chromatin has a role in the transcription regulation of genes important for melanoma initiation and progression.

**Figure 8.**
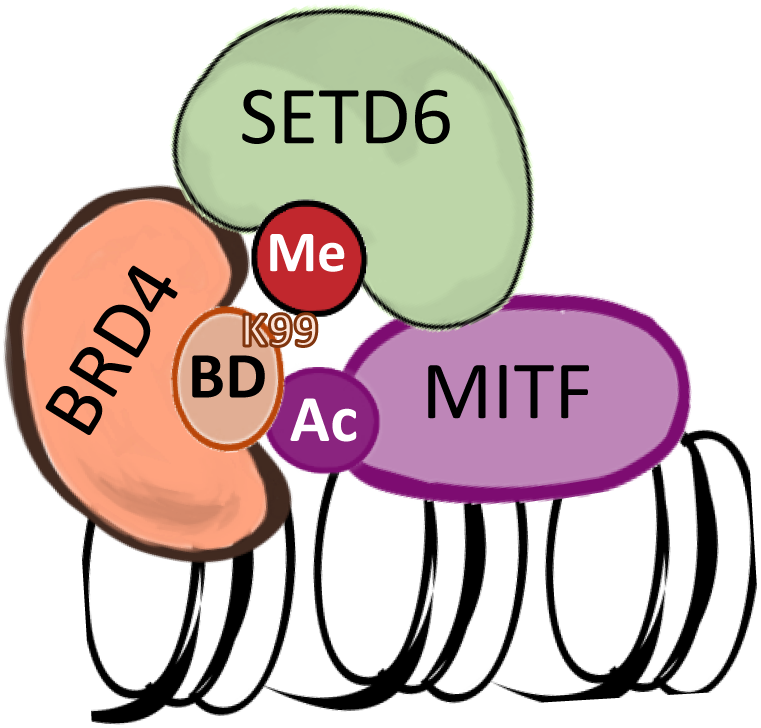
BRD4 interacts with acetylated MITF by its BD in a SETD6 dependent manner. Schematic model illustrating the proposed SETD6-BRD4-MITF axis

## Discussion

The PKMT SETD6 has a fundamental role in the regulation of several biological processes and pathologies including cancer (17–22,24–30). The tight and significant correlation between the high SETD6 expression level and poor survival of melanoma patients led us to study its role in this cancer. RNA-Sequencing revealed that SETD6 regulates the expression of genes which are related to various pathways in melanoma. Accordingly, alterations in cellular phenotypes, such as cell proliferation and adhesion, were seen after SETD6 knockout, providing the first evidence of its involvement in melanoma. These phenotypes are also consistent with previous studies in human cancers such as breast, glioma and others (18,27,28,30). The increasing cellular functions of SETD6 both in normal and disease states, make it an attractive candidate for therapeutic use.

We have previously shown that SETD6 specifically methylates BRD4 at K99 in a breast cancer cellular model, which in turn regulates the transcription of genes that control mRNA translation (28). Here we provide evidence that BRD4 methylation by SETD6 at K99 plays an important role also in melanoma cells. Our biochemical and genomics analysis revealed that SETD6 mediated methylation of BRD4 at chromatin is essential to drive selective gene expression programs by recruitment of BRD4 to different genomic loci and genes. This effect is both SETD6 and BRD4 K99 methylation dependent. Mechanistically we observed two independent modes of action in breast cancer vs. melanoma cells. In breast cancer cells BRD4 methylation specifically determines the recruitment of the transcription factor E2F1 to selected target genes. However, here we provide evidence that in melanoma cells the methylation of BRD4 is required for the recruitment of MITF. It seems that methylated BRD4 at chromatin might serve as a scaffold for recruiting different transcription factors to allow selective and efficient regulation of gene expression programs.

Our genomic analysis revealed that MITF genomic occupancy is regulated by SETD6, similar to BRD4. Interestingly, our study also uncovered that SETD6 physically interacts with MITF. This observation prompted us to investigate whether there is an overlap between the target genes of BRD4 and MITF that are regulated by the presence of SETD6. Indeed, we observed common genomic loci for BRD4 and MITF that are affected by SETD6. Consistent with our findings, a recent study showed that BRD4 and MITF physically interact with each other in melanocyte cells and this interaction controls the production of melanin(2). These observations led us to examine if BRD4 and MITF co-occupy the same genomic regions. Our data revealed that this co-occupancy also depends on SETD6 and the methylation of BRD4 at K99. Future work will reveal if SETD6 and the methylation of BRD4 are also required for BRD4 and MITF transcriptional activity in regulating specific gene expression programs in melanoma.

Previously, MITF acetylation by p300 was shown to direct MITF to distinct genomic locations, thereby coordinating the expression of genes which regulate melanocyte and melanoma proliferation and differentiation (56,57). This led us to investigate whether the newly identified BRD4-MITF interaction in melanoma cells is mediated by the bromo domain of BRD4 and acetylated MITF. We tackled this question by either inhibiting the bromo domain of BRD4 or inducing MITF acetylation in cells. Our findings not only support this model, but also suggest that it is regulated by SETD6 and the methylation of BRD4 at K99. A particularly intriguing direction for future research is to explore the direct interaction between MITF and SETD6. Further investigation into the specific acetylation sites in MITF facilitating this interaction, as well as the potential crosstalk with nearby post-translational modifications on MITF, would be of significant interest as well.

In summary, we uncovered a new functional crosstalk at chromatin, between SETD6, BRD4, and MITF. The formation of this chromatin-associated complex plays a critical role in selective recruitment of these TFs to different genomic loci in melanoma cells. Thus, targeting the biochemical interactions identified in this study might represent a promising strategy for inhibiting melanoma progression. The inhibition of BRD4 methylation by SETD6 or preventing BRD4 from recognizing MITF acetylation may yield similar therapeutic outcomes.

## Acknowledgments

We thank the Levy and Bernstein labs for technical assistance and helpful discussions. This work was supported by grants to DL from The Israel Science Foundation (262/18 and 496/23), The Research Career Development Award from the Israel Cancer Research Fund and from the Israel Cancer Association. DL and EB are supported by a US-Israel Binational Science Foundation grant.

## Author Contribution

TEB and DL conceived and designed the majority of the experiments. NTM and LL performed the bioinformatics analysis. DS assisted in the microscopic data analysis. MF, TD, EB and CG helped with experimental design and provided valuable conceptual input for the study. TEB and DL wrote the paper. EB and DL acquired funding. All authors read and approved the final manuscript.

---

**Supplementary Fig 1.**
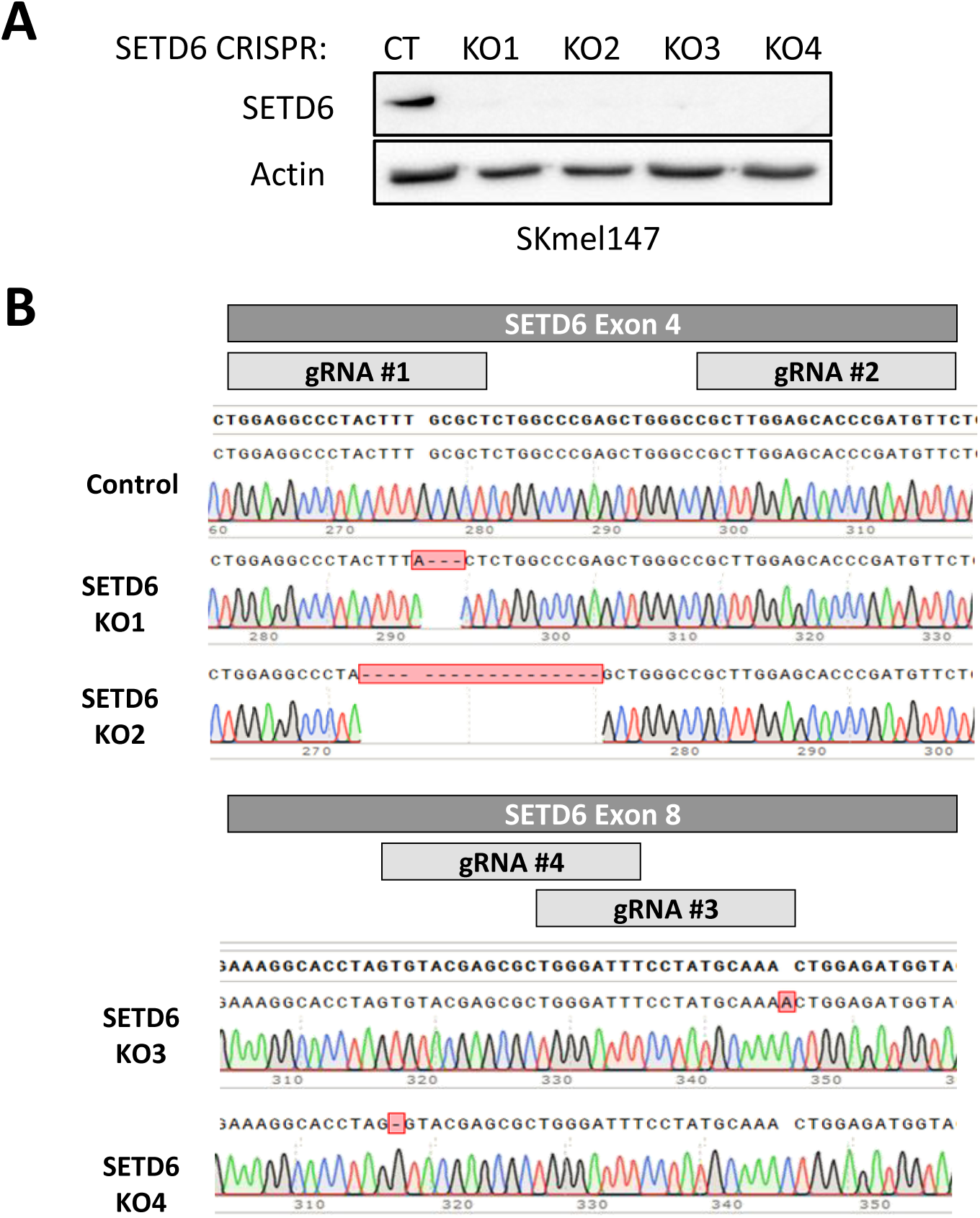

**Supplementary Fig 2.**
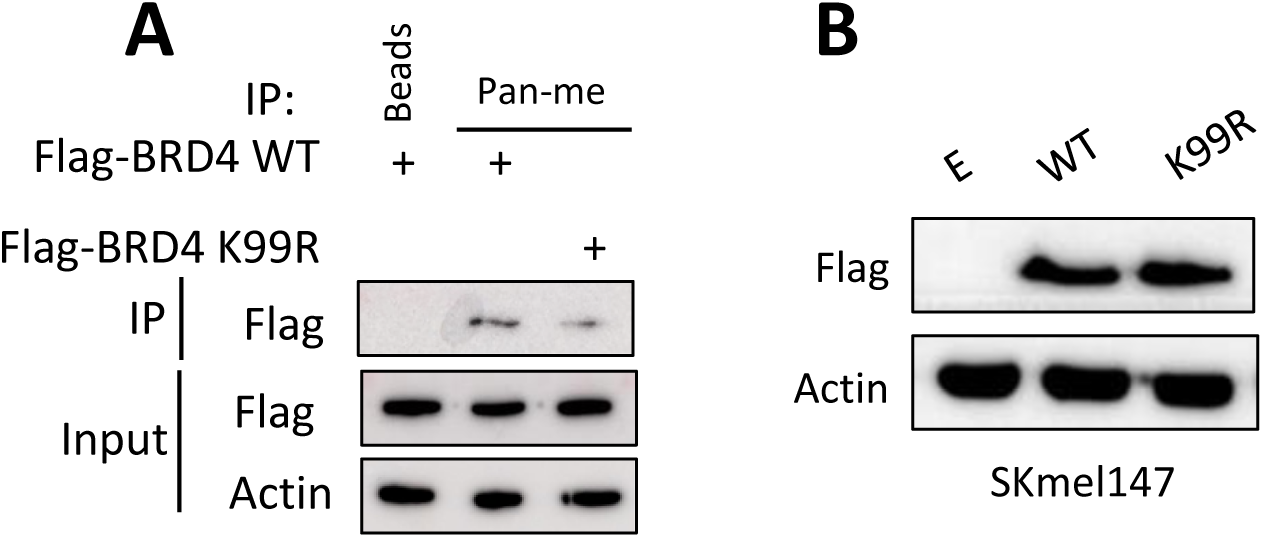

**Supplementary Fig 3.**
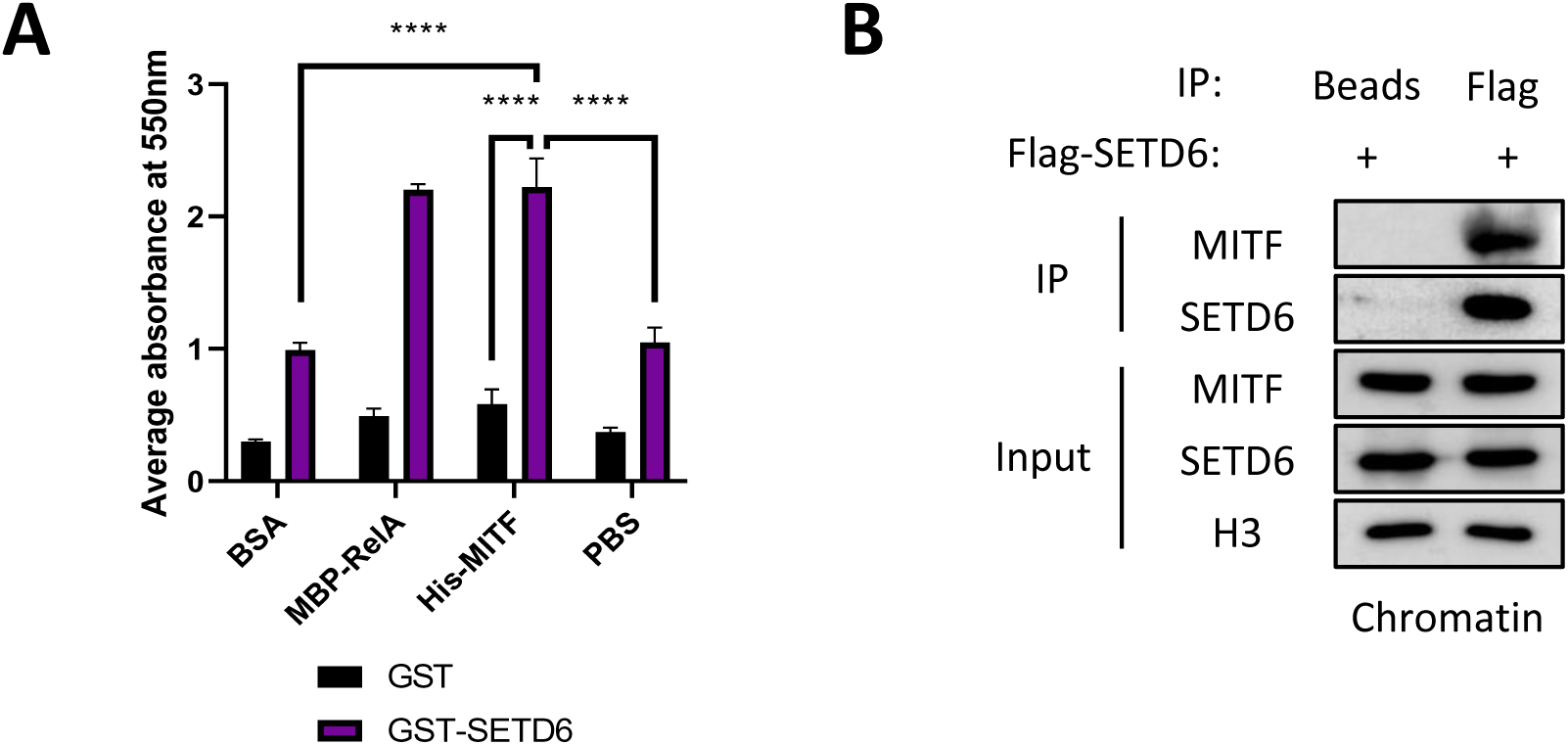

**Supplementary Fig 4.**
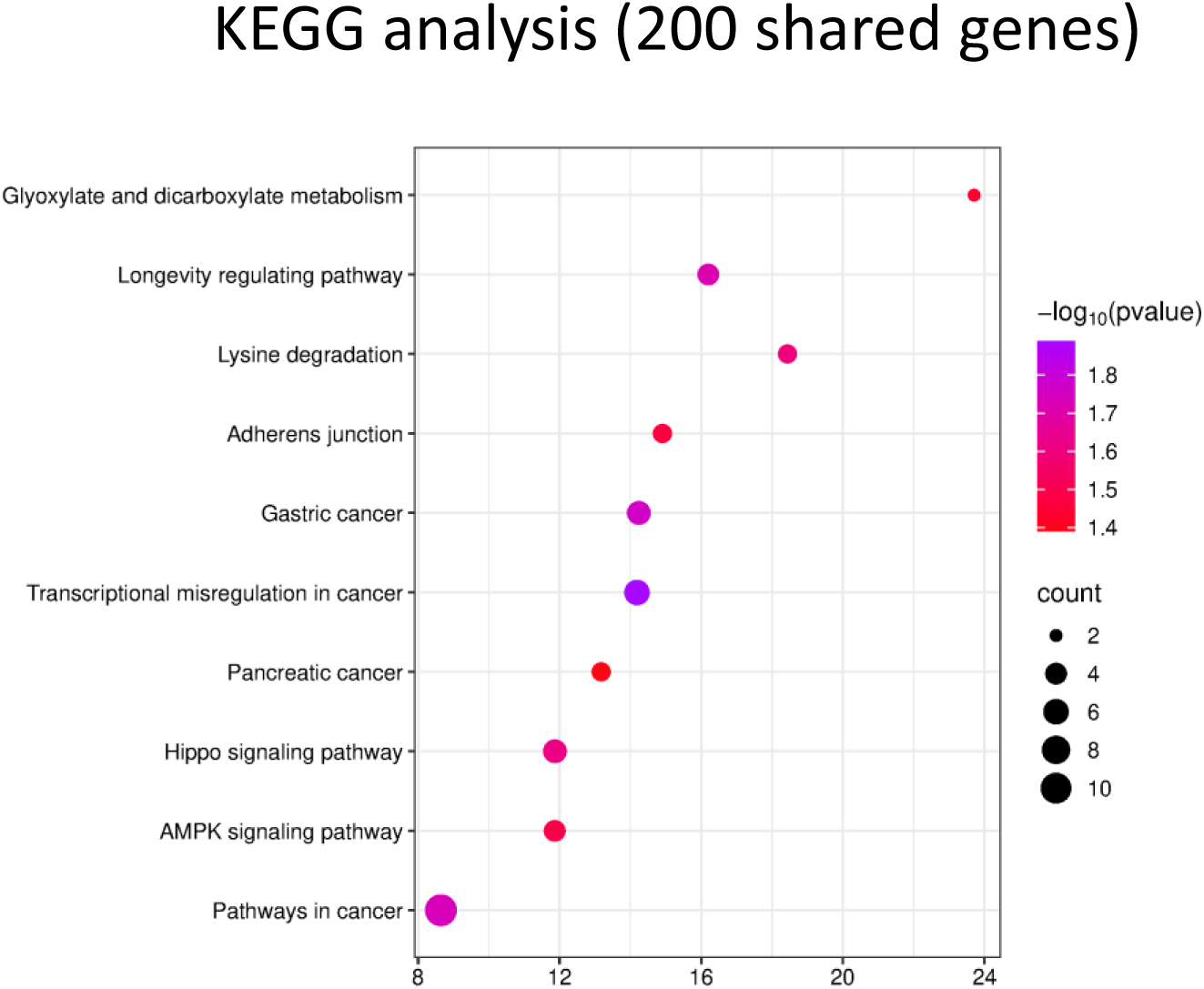

**Supplementary Fig 5.**
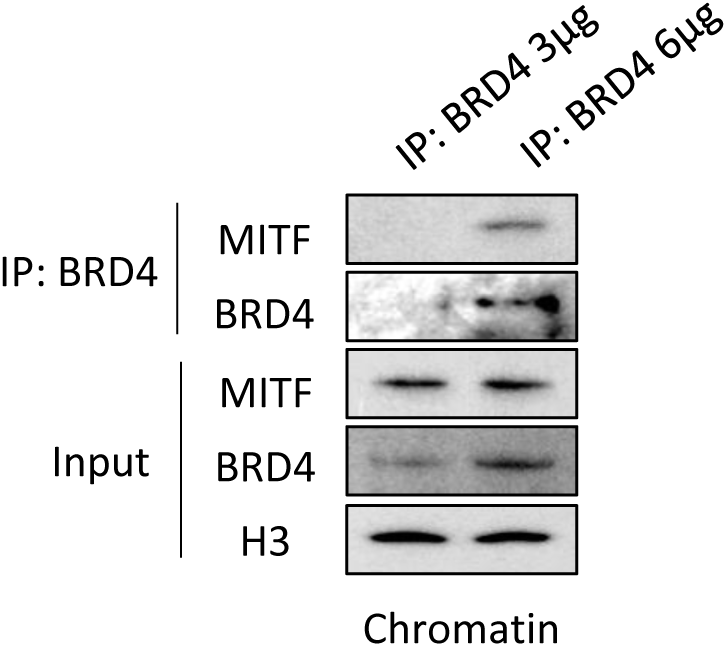

**Supplementary Fig 6.**
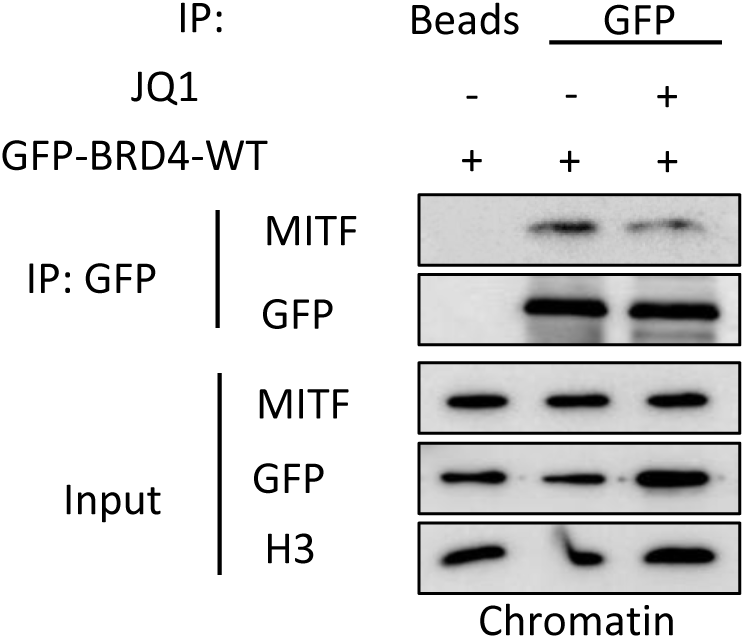

**Supplementary Table 1.**
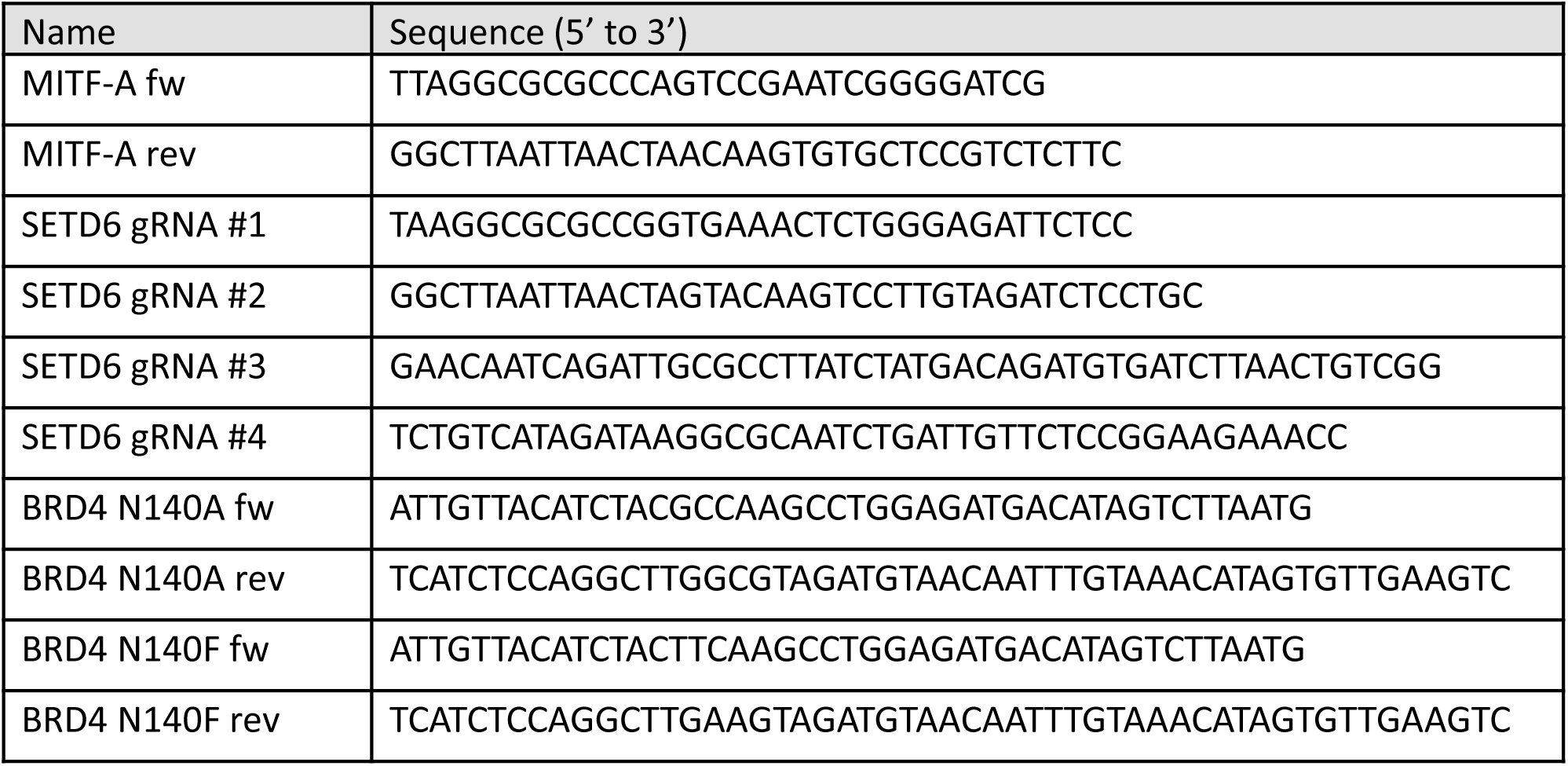

## References

1. Levy, C., Khaled, M. and Fisher, D.E. (2006) MITF: master regulator of melanocyte development and melanoma oncogene. Trends Mol Med, 12, 406–414.

2. Sobiepanek, A., Paone, A., Cutruzzola, F. and Kobiela, T. (2021) Biophysical characterization of melanoma cell phenotype markers during metastatic progression. Eur Biophys J, 50, 523–542.

3. Sarkar, D., Leung, E.Y., Baguley, B.C., Finlay, G.J. and Askarian-Amiri, M.E. (2015) Epigenetic regulation in human melanoma: past and future. Epigenetics, 10, 103–121.

4. Goding, C.R. and Arnheiter, H. (2019) MITF-the first 25 years. Genes Dev, 33, 983–1007.

5. Garraway, L.A., Widlund, H.R., Rubin, M.A., Getz, G., Berger, A.J., Ramaswamy, S., Beroukhim, R., Milner, D.A., Granter, S.R., Du, J. et al. (2005) Integrative genomic analyses identify MITF as a lineage survival oncogene amplified in malignant melanoma. Nature, 436, 117–122.

6. Filippakopoulos, P. and Knapp, S. (2014) Targeting bromodomains: epigenetic readers of lysine acetylation. Nat Rev Drug Discov, 13, 337–356.

7. Morgado-Pascual, J.L., Rayego-Mateos, S., Tejedor, L., Suarez-Alvarez, B. and Ruiz-Ortega, M. (2019) Bromodomain and Extraterminal Proteins as Novel Epigenetic Targets for Renal Diseases. Front Pharmacol, 10, 1315.

8. Delmore, J.E., Issa, G.C., Lemieux, M.E., Rahl, P.B., Shi, J., Jacobs, H.M., Kastritis, E., Gilpatrick, T., Paranal, R.M., Qi, J. et al. (2011) BET bromodomain inhibition as a therapeutic strategy to target c-Myc. Cell, 146, 904–917.

9. Shi, J., Wang, Y., Zeng, L., Wu, Y., Deng, J., Zhang, Q., Lin, Y., Li, J., Kang, T., Tao, M. et al. (2014) Disrupting the interaction of BRD4 with diacetylated Twist suppresses tumorigenesis in basal-like breast cancer. Cancer Cell, 25, 210–225.

10. Segura, M.F., Fontanals-Cirera, B., Gaziel-Sovran, A., Guijarro, M.V., Hanniford, D., Zhang, G., Gonzalez-Gomez, P., Morante, M., Jubierre, L., Zhang, W. et al. (2013) BRD4 sustains melanoma proliferation and represents a new target for epigenetic therapy. Cancer Res, 73, 6264–6276.

11. Trivedi, A., Mehrotra, A., Baum, C.E., Lewis, B., Basuroy, T., Blomquist, T., Trumbly, R., Filipp, F.V., Setaluri, V. and de la Serna, I.L. (2020) Bromodomain and extra-terminal domain (BET) proteins regulate melanocyte differentiation. Epigenetics Chromatin, 13, 14.

12. Deribe, Y.L., Pawson, T. and Dikic, I. (2010) Post-translational modifications in signal integration. Nat Struct Mol Biol, 17, 666–672.

13. Alam, H., Gu, B. and Lee, M.G. (2015) Histone methylation modifiers in cellular signaling pathways. Cell Mol Life Sci, 72, 4577–4592.

14. Hamamoto, R., Saloura, V. and Nakamura, Y. (2015) Critical roles of non-histone protein lysine methylation in human tumorigenesis. Nat Rev Cancer, 15, 110–124.

15. Levy, D. (2019) Lysine methylation signaling of non-histone proteins in the nucleus. Cell Mol Life Sci, 76, 2873–2883.

16. Kouzarides, T. (2007) Chromatin modifications and their function. Cell, 128, 693–705.

17. Arrowsmith, C.H., Bountra, C., Fish, P.V., Lee, K. and Schapira, M. (2012) Epigenetic protein families: a new frontier for drug discovery. Nat Rev Drug Discov, 11, 384–400.

18. O’Neill, D.J., Williamson, S.C., Alkharaif, D., Monteiro, I.C., Goudreault, M., Gaughan, L., Robson, C.N., Gingras, A.C. and Binda, O. (2014) SETD6 controls the expression of estrogen-responsive genes and proliferation of breast carcinoma cells. Epigenetics, 9, 942–950.

19. Admoni-Elisha, L., Elbaz, T., Chopra, A., Shapira, G., Bedford, M.T., Fry, C.J., Shomron, N., Biggar, K., Feldman, M. and Levy, D. (2022) TWIST1 methylation by SETD6 selectively antagonizes LINC-PINT expression in glioma. Nucleic Acids Res, 50, 6903–6918.

20. Binda, O., Sevilla, A., LeRoy, G., Lemischka, I.R., Garcia, B.A. and Richard, S. (2013) SETD6 monomethylates H2AZ on lysine 7 and is required for the maintenance of embryonic stem cell self-renewal. Epigenetics, 8, 177–183.

21. Chen, A., Feldman, M., Vershinin, Z. and Levy, D. (2016) SETD6 is a negative regulator of oxidative stress response. Biochim Biophys Acta, 1859, 420–427.

22. Feldman, M., Vershinin, Z., Goliand, I., Elia, N. and Levy, D. (2019) The methyltransferase SETD6 regulates Mitotic progression through PLK1 methylation. Proc Natl Acad Sci U S A, 116, 1235–1240.

23. Kublanovsky, M., Ulu, G.T., Weirich, S., Levy, N., Feldman, M., Jeltsch, A. and Levy, D. (2023) Methylation of the transcription factor E2F1 by SETD6 regulates SETD6 expression via a positive feedback mechanism. J Biol Chem, 299, 105236.

24. Levy, D., Kuo, A.J., Chang, Y., Schaefer, U., Kitson, C., Cheung, P., Espejo, A., Zee, B.M., Liu, C.L., Tangsombatvisit, S. et al. (2011) Lysine methylation of the NF-kappaB subunit RelA by SETD6 couples activity of the histone methyltransferase GLP at chromatin to tonic repression of NF-kappaB signaling. Nat Immunol, 12, 29–36.

25. Martin-Morales, L., Feldman, M., Vershinin, Z., Garre, P., Caldes, T. and Levy, D. (2017) SETD6 dominant negative mutation in familial colorectal cancer type X. Hum Mol Genet, 26, 4481–4493.

26. Vershinin, Z., Feldman, M., Chen, A. and Levy, D. (2016) PAK4 Methylation by SETD6 Promotes the Activation of the Wnt/beta-Catenin Pathway. J Biol Chem, 291, 6786–6795.

27. Vershinin, Z., Feldman, M. and Levy, D. (2020) PAK4 methylation by the methyltransferase SETD6 attenuates cell adhesion. Sci Rep, 10, 17068.

28. Vershinin, Z., Feldman, M., Werner, T., Weil, L.E., Kublanovsky, M., Abaev-Schneiderman, E., Sklarz, M., Lam, E.Y.N., Alasad, K., Picaud, S. et al. (2021) BRD4 methylation by the methyltransferase SETD6 regulates selective transcription to control mRNA translation. Sci Adv, 7.

29. Weil, L.E., Shmidov, Y., Kublanovsky, M., Morgenstern, D., Feldman, M., Bitton, R. and Levy, D. (2018) Oligomerization and Auto-methylation of the Human Lysine Methyltransferase SETD6. J Mol Biol, 430, 4359–4368.

30. Admoni-Elisha, L., Abaev-Schneiderman, E., Cohn, O., Shapira, G., Shomron, N., Feldman, M. and Levy, D. (2022) Structure-function conservation between the methyltransferases SETD3 and SETD6. Biochimie, 200, 27–35.

31. Li, B. and Dewey, C.N. (2011) RSEM: accurate transcript quantification from RNA-Seq data with or without a reference genome. BMC Bioinformatics, 12, 323.

32. Ewels, P., Magnusson, M., Lundin, S. and Kaller, M. (2016) MultiQC: summarize analysis results for multiple tools and samples in a single report. Bioinformatics, 32, 3047–3048.

33. Love, M.I., Huber, W. and Anders, S. (2014) Moderated estimation of fold change and dispersion for RNA-seq data with DESeq2. Genome Biol, 15, 550.

35. Lachmann, A., Xu, H., Krishnan, J., Berger, S.I., Mazloom, A.R. and Ma’ayan, A. (2010) ChEA: transcription factor regulation inferred from integrating genome-wide ChIP-X experiments. Bioinformatics, 26, 2438–2444.

35. (2018) NeatSeq-Flow: A Lightweight High-Throughput Sequencing Workflow Platform for Non-Programmers and Programmers Alike. bioRxiv.

36. Li, H. and Durbin, R. (2009) Fast and accurate short read alignment with Burrows-Wheeler transform. Bioinformatics, 25, 1754–1760.

37. Danecek, P., Bonfield, J.K., Liddle, J., Marshall, J., Ohan, V., Pollard, M.O., Whitwham, A., Keane, T., McCarthy, S.A., Davies, R.M. et al. (2021) Twelve years of SAMtools and BCFtools. Gigascience, 10.

38. Ramirez, F., Ryan, D.P., Gruning, B., Bhardwaj, V., Kilpert, F., Richter, A.S., Heyne, S., Dundar, F. and Manke, T. (2016) deepTools2: a next generation web server for deep-sequencing data analysis. Nucleic Acids Res, 44, W160–165.

39. Amemiya, H.M., Kundaje, A. and Boyle, A.P. (2019) The ENCODE Blacklist: Identification of Problematic Regions of the Genome. Sci Rep, 9, 9354.

40. Quinlan, A.R. and Hall, I.M. (2010) BEDTools: a flexible suite of utilities for comparing genomic features. Bioinformatics, 26, 841–842.

41. Zhang, Y., Liu, T., Meyer, C.A., Eeckhoute, J., Johnson, D.S., Bernstein, B.E., Nusbaum, C., Myers, R.M., Brown, M., Li, W. et al. (2008) Model-based analysis of ChIP-Seq (MACS). Genome Biol, 9, R137.

42. Meers, M.P., Tenenbaum, D. and Henikoff, S. (2019) Peak calling by Sparse Enrichment Analysis for CUT&RUN chromatin profiling. Epigenetics Chromatin, 12, 42.

43. Robinson, J.T., Thorvaldsdottir, H., Winckler, W., Guttman, M., Lander, E.S., Getz, G. and Mesirov, J.P. (2011) Integrative genomics viewer. Nat Biotechnol, 29, 24–26.

44. Wang, Q., Li, M., Wu, T., Zhan, L., Li, L., Chen, M., Xie, W., Xie, Z., Hu, E., Xu, S., et al. (2022) Exploring Epigenomic Datasets by ChIPseeker. Curr Protoc, 2, e585.

45. Yu, G., Wang, L.G. and He, Q.Y. (2015) ChIPseeker: an R/Bioconductor package for ChIP peak annotation, comparison and visualization. Bioinformatics, 31, 2382–2383.

46. Schindelin, J., Arganda-Carreras, I., Frise, E., Kaynig, V., Longair, M., Pietzsch, T., Preibisch, S., Rueden, C., Saalfeld, S., Schmid, B. et al. (2012) Fiji: an open-source platform for biological-image analysis. Nature methods, 9, 676–682.

47. Legland, D., Arganda-Carreras, I. and Andrey, P. (2016) MorphoLibJ: integrated library and plugins for mathematical morphology with ImageJ. Bioinformatics, 32, 3532–3534.

48. Dissanayake, S.K., Wade, M., Johnson, C.E., O’Connell, M.P., Leotlela, P.D., French, A.D., Shah, K.V., Hewitt, K.J., Rosenthal, D.T., Indig, F.E. et al. (2007) The Wnt5A/protein kinase C pathway mediates motility in melanoma cells via the inhibition of metastasis suppressors and initiation of an epithelial to mesenchymal transition. J Biol Chem, 282, 17259–17271.

49. Gajos-Michniewicz, A. and Czyz, M. (2020) WNT Signaling in Melanoma. Int J Mol Sci, 21.

50. Mahabeleshwar, G.H. and Byzova, T.V. (2007) Angiogenesis in melanoma. Semin Oncol, 34, 555–565.

51. Skene, P.J. and Henikoff, S. (2017) An efficient targeted nuclease strategy for high-resolution mapping of DNA binding sites. Elife, 6.

52. Chen, E.Y., Tan, C.M., Kou, Y., Duan, Q., Wang, Z., Meirelles, G.V., Clark, N.R. and Ma’ayan, A. (2013) Enrichr: interactive and collaborative HTML5 gene list enrichment analysis tool. BMC Bioinformatics, 14, 128.

53. Huang, B., Yang, X.D., Zhou, M.M., Ozato, K. and Chen, L.F. (2009) Brd4 coactivates transcriptional activation of NF-kappaB via specific binding to acetylated RelA. Mol Cell Biol, 29, 1375–1387.

54. Xie, F., Xiao, X., Tao, D., Huang, C., Wang, L., Liu, F., Zhang, H., Niu, H. and Jiang, G. (2020) circNR3C1 Suppresses Bladder Cancer Progression through Acting as an Endogenous Blocker of BRD4/C-myc Complex. Mol Ther Nucleic Acids, 22, 510–519.

55. Yu, J., Chen, M., Sang, Q., Li, F., Xu, Z., Yu, B., He, C., Su, L., Dai, W., Yan, C. et al. (2024) Super-enhancer Activates Master Transcription Factor NR3C1 Expression and Promotes 5-FU Resistance in Gastric Cancer. Adv Sci (Weinh), e2409050.

56. Louphrasitthiphol, P., Loffreda, A., Pogenberg, V., Picaud, S., Schepsky, A., Friedrichsen, H., Zeng, Z., Lashgari, A., Thomas, B., Patton, E.E. et al. (2023) Acetylation reprograms MITF target selectivity and residence time. Nat Commun, 14, 6051.

57. Louphrasitthiphol, P., Siddaway, R., Loffreda, A., Pogenberg, V., Friedrichsen, H., Schepsky, A., Zeng, Z., Lu, M., Strub, T., Freter, R. et al. (2020) Tuning Transcription Factor Availability through Acetylation-Mediated Genomic Redistribution. Mol Cell, 79, 472–487 e410.

58. Butler, L.M., Zhou, X., Xu, W.S., Scher, H.I., Rifkind, R.A., Marks, P.A. and Richon, V.M. (2002) The histone deacetylase inhibitor SAHA arrests cancer cell growth, up-regulates thioredoxin-binding protein-2, and down-regulates thioredoxin. Proc Natl Acad Sci U S A, 99, 11700–11705.

59. Filippakopoulos, P., Qi, J., Picaud, S., Shen, Y., Smith, W.B., Fedorov, O., Morse, E.M., Keates, T., Hickman, T.T., Felletar, I. et al. (2010) Selective inhibition of BET bromodomains. Nature, 468, 1067–1073.

60. Jung, M., Philpott, M., Muller, S., Schulze, J., Badock, V., Eberspacher, U., Moosmayer, D., Bader, B., Schmees, N., Fernandez-Montalvan, A. et al. (2014) Affinity map of bromodomain protein 4 (BRD4) interactions with the histone H4 tail and the small molecule inhibitor JQ1. J Biol Chem, 289, 9304–9319.

